# Core circadian clock transcription factor BMAL1 regulates mammary epithelial cell growth, differentiation, and milk component synthesis

**DOI:** 10.1101/2021.02.23.432439

**Authors:** Theresa Casey, Aridany Suarez-Trujillo, Shelby Cummings, Katelyn Huff, Jennifer Crodian, Ketaki Bhide, Clare Aduwari, Kelsey Teeple, Avi Shamay, Sameer J. Mabjeesh, Phillip San Miguel, Jyothi Thimmapuram, Karen Plaut

**Affiliations:** Department of Animal Science, Purdue University, West Lafayette, IN, USA; Bioinformatics Core, Purdue University; Animal Science Institute, Agriculture Research Origination, The Volcani Center, Rishon Letsiyon, Israel; Department of Animal Sciences, The Robert H. Smith Faculty of Agriculture, Food, and Environment, The Hebrew University of Jerusalem, Rehovot, Israel; Genomics Core, Purdue University

**Author notes:** **Address for correspondence.** Theresa M. Casey, BCHM Room 326, 175 South University St., West Lafayette, IN 47907.

## Abstract

The role the mammary epithelial circadian clock plays in gland development and lactation is unknown. We hypothesized that mammary epithelial clocks function to regulate mammogenesis and lactogenesis, and propose the core clock transcription factor BMAL1:CLOCK regulates genes that control mammary epithelial development and milk synthesis. Our objective was to identify transcriptional targets of BMAL1 in undifferentiated (UNDIFF) and lactogen differentiated (DIFF) mammary epithelial cells (HC11) using ChIP-seq. Ensembl gene IDs with the nearest transcriptional start site to peaks were explored as potential targets, and represented 846 protein coding genes common to UNDIFF and DIFF cells and 2773 unique to DIFF samples. Genes with overlapping peaks between samples (1343) enriched cell-cell adhesion, membrane transporters and lipid metabolism categories. To functionally verify targets, an HC11 line with *Bmal1* gene knocked out (BMAL1-KO) using CRISPR-CAS was created. BMAL1-KO cultures had lower cell densities over an eight-day growth curve, which was associated with increased (p<0.05) levels of reactive oxygen species and lower expression of superoxide dismutase 3 (*Sod3*). Q-PCR analysis also found lower expression of the putative targets, prolactin receptor (*Prlr*), *Ppara*, and beta-casein (*Csn2*). Findings support our hypothesis and highlight potential importance of clock in mammary development and substrate transport.

## INTRODUCTION

Circadian clocks set daily rhythms of gene expression to maintain tissue homeostasis and coordinate cellular metabolism[1–3]. In mammary tissue, temporal expression patterns of core clock genes change across reproductive states of the female, and appear to be associated with changes in developmental stage of the gland [4, 5]. However, the significance of these changes and the role of the molecular clock and core clock genes’ functions in mammary development and lactation is currently unknown.

Clocks’ molecular mechanism generate circadian rhythms through a series of interlocked transcription-translation feedback loops. CLOCK and BMAL1 are at the core of the loop, and as a heterodimer (BMAL1:CLOCK) function as a transcription factor that binds the enhancer box (E-box) regulatory element in promoter regions of genes[6]. *Period* (*Per1, Per2* and *Per3*) and *Cryptochrome* (*Cry1* and *Cry2*) genes are transcriptional targets of BMAL1:CLOCK and form the negative arm of the core circadian clock loop, whereby PER and CRY proteins together inhibit CLOCK:BMAL1-mediated transcription[7–10]. The 24 hr periodicity in activation-repression of elements of the core molecular clock results in corresponding circadian rhythms of gene expression in about 10% of the transcriptome[8].

Although there is some overlap, the rhythmic output genes of clocks differ among tissues, allowing circadian control of function and activity appropriate for each organ. Circadian and cell cycle regulation is coupled to coordinate cell proliferation with tissue function across multiple organs[8, 11, 12]. This coupling likely also exists in the mammary gland, as temporal analysis of mammary transcriptomes of mature virgin mice[13] and lactating women[14] demonstrated genes that regulate cell growth and differentiation exhibit circadian rhythms of expression. Moreover, the *Clock-Δ19* mice, which have a point mutation that results in down regulation of CLOCK-BMAL1 target genes, exhibit decreased lactation competence as evident in lower rates of pup growth and increased rates of neonatal mortality[15–17]. The decreased postnatal pup survival was associated with impaired mammary development in *Clock-Δ19* dams[18], and isolated mammary epithelial cells from virgin *Clock-Δ19* mice had lower stem cell-like properties[13]. Whereas reduction of CLOCK protein in a mammary epithelial cell line with shRNA affected cell growth and decreased expression of factors associated with mammary differentiation and milk synthesis[18].

With this knowledge, we hypothesized that circadian clocks in the mammary gland play an integral role in regulating mammogenesis. In particular, we proposed that the transcription factor BMAL1:CLOCK functions to regulate expression of genes that control mammary epithelial development. Moreover, the observation that stoichiometry changed between positive (BMAL:CLOCK) and negative (PER2) elements of core circadian clock components in the mammary gland during the transition from pregnancy to lactation [5] led us to the hypothesize that transcriptional targets of BMAL1:CLOCK change as the gland differentiates and function to regulate milk component synthesis. The objectives of this study were to use ChIP-seq analysis to identify transcriptional targets of BMAL1, and determine if targets of BMAL1 changed upon differentiation of mammary epithelial cells. A line of mammary epithelial cells with the BMAL1 gene knocked out (BMAL1-KO) using CRISPR-CAS was created to verify ChIP-seq findings, and study functional effects of loss of gene on growth and differentiation of cells in culture.

## RESULTS

### Validation of BMAL1 antibody and quality of ChIP-seq data

To test the hypothesis that BMAL1 transcriptional targets changed with state of mammary epithelial differentiation we used ChIP-seq analysis to identify genes potentially regulated by BMAL1 in undifferentiated (UNDIFF) and lactogen hormone differentiated (DIFF) HC11 cultures. Prior to beginning studies, specificity of the ChIP-grade antibody for the BMAL1 protein was confirmed with immunoprecipitation and western blot analysis (Supplemental Figure S1a). Nanochip analysis of sonicated input DNA indicated that optimal size of sheared fragments was achieved for ChIP-seq analysis (Supplemental Figure S1b). Evaluation of antibody specificity indicated no difference between mock-ChIP and BMAL1-ChIP samples in the cycle threshold values following q-PCR analysis of an exon region of the *Magea1_2* sperm specific gene, which is not a BMAL1:CLOCK target. Whereas a 9-fold difference in enrichment was found between q-PCR product of BMAL1-ChIP and mock-ChIP for the *Per1* promoter region versus the exon region of *Magea1_2* (Supplemental Figure S1c).

The eight samples sequenced for these studies consisted of four pairs of input and ChIP samples with two pairs from UNDIFF and two from DIFF HC11 cultures. Across the four samples the amount of DNA captured by ChIP was 1.03 ± 0.24% of the input DNA. Mapping rate of reads to the mouse genome across the four paired input-ChIP samples averaged 96% (Supplemental Table S1). For UNDIFF samples, there was an average of ∼6,000 peaks with a 2-fold enrichment over input tag count (FDR ≤ 0.01), and over 13,000 peaks were identified across both DIFF samples. Annotation analysis of peaks found similar frequencies of location within the genome across the samples, with approximately 62% of peaks located in intergenic sites and 34% within introns (Figure 1a). Less than 1% of peaks were found in 3’ untranslated region, transcriptional termination sites (TSS), and within exons.

**Figure 1.**
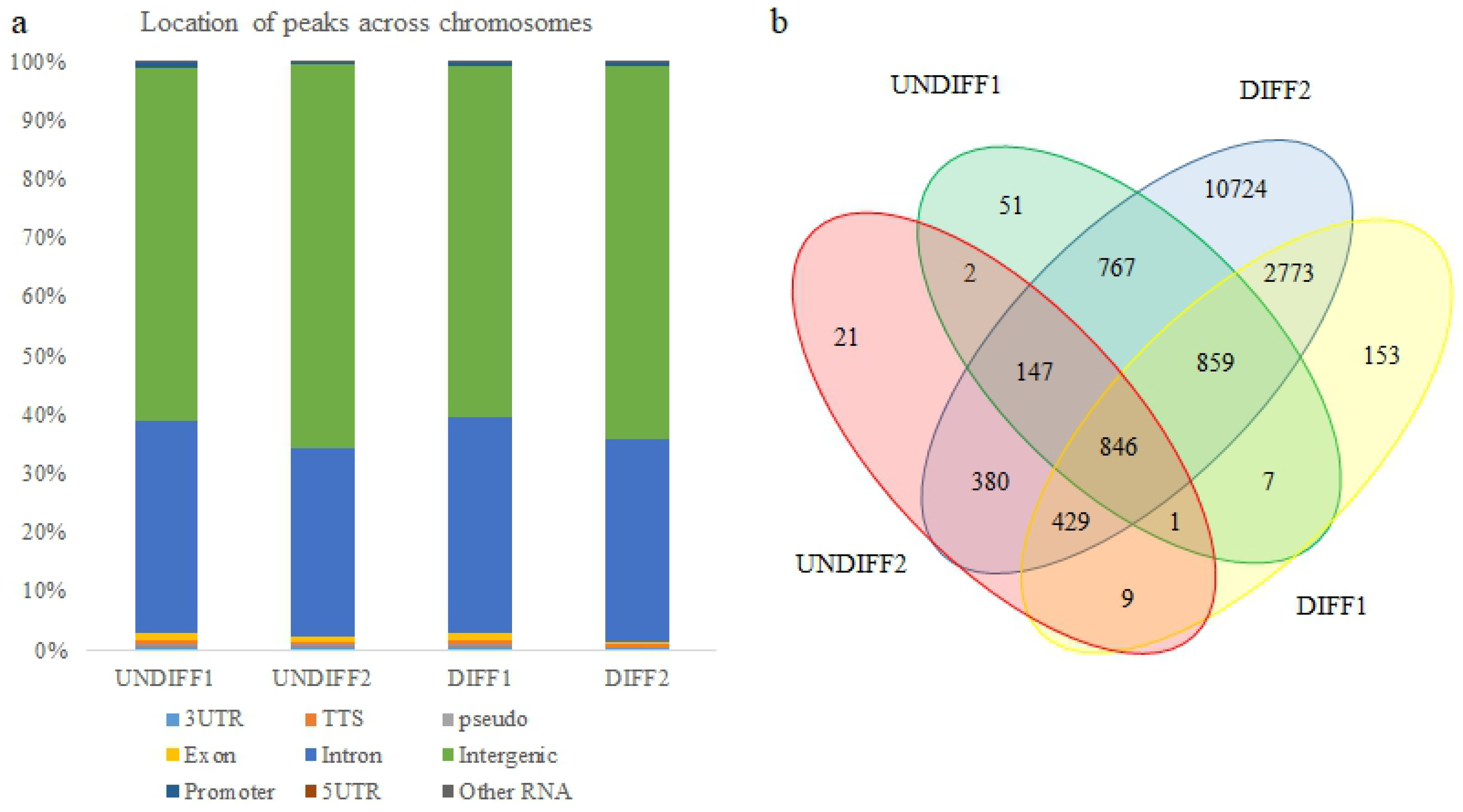
(a) Homer annotation analysis of BMAL1 ChIP-seq peak location shows frequency of distribution across the genome in undifferentiated (UNDIFF) and lactogen differentiated (DIFF) HC11 cultures. 3UTR, 3’ untranslated region; TTS, terminal transcription site; pseudo, pseudogene; 5UTR, 5’ untranslated region. (b) Venn diagram illustrating the overlap and number of unique Ensembl gene IDs of protein coding genes with transcriptional start site nearest to BMAL1 ChIP-seq peaks in the two undifferentiated (UNDIFF1 and UNDIFF2) and two differentiated (DIFF1 and DIFF2) samples.

**Table 1.**
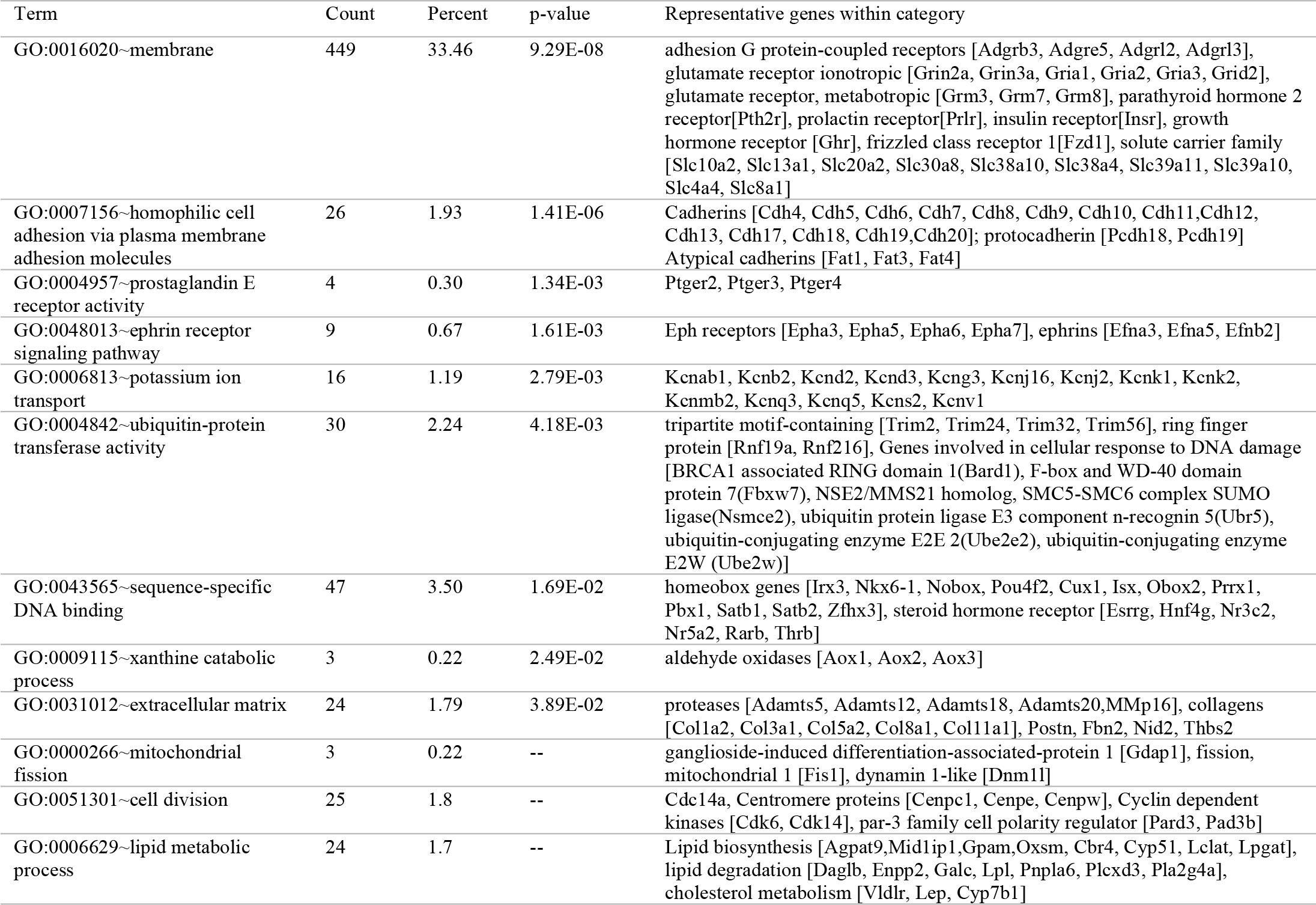
Representative categories enriched with protein coding genes nearest to BMAL1 ChIP-seq overlapping peaks

### Functional analysis of protein coding genes nearest transcriptional start sites of ChIP-seq peaks

Ensembl protein coding gene IDs with the nearest transcriptional start site to peaks were explored as potential regulatory targets of BMAL1. Analysis of overlapping gene IDs among the samples found 846 common to all four ChIPed samples (i.e. UNDIFF and DIFF; Figure 1b; Supplemental Table S2). There were 2773 protein coding genes common to both DIFF samples, but not found in UNDIFF samples, and were considered as potential BMAL1 targets distinctly regulated in differentiated versus undifferentiated mammary epithelial cells (Supplemental Table S3). Potential BMAL1 target genes with overlapping peaks between at least two samples were considered high confidence and used for downstream analysis. There were 97 common between UNDIFF samples (Supplemental Table S4) and 778 common between DIFF samples (Supplemental Table S5). Gene targets with overlapping peaks between any two samples (1343; Supplemental Table S6), regardless of differentiation state, were used for downstream analysis.

**Table 2.**
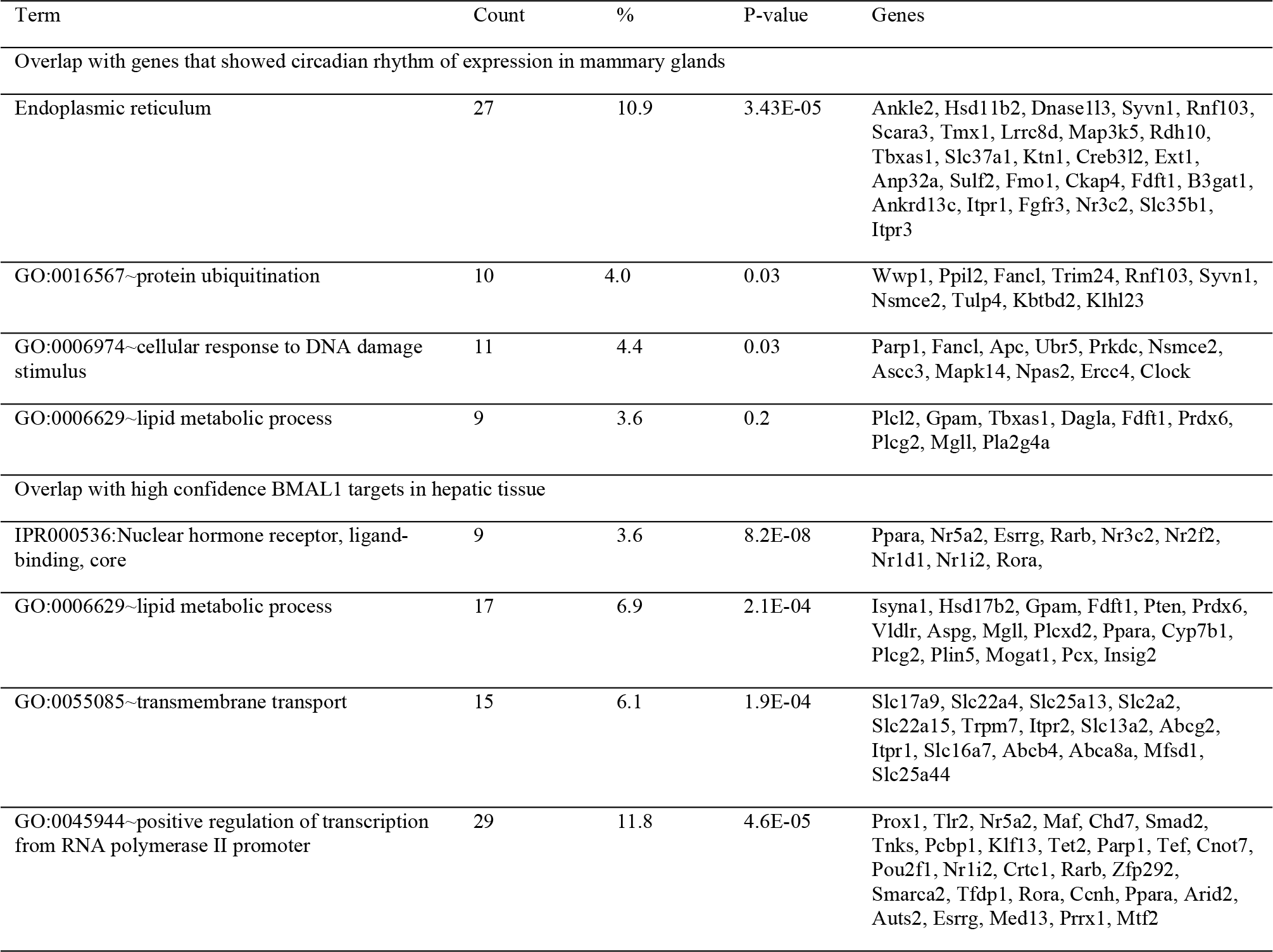
Representative categories enriched with potential BMAL1 target genes identified using ChIP-seq that overlapped with genes that showed circadian rhythms of expression in mammary glands and high confidence BMAL1 targets identified in hepatic tissue

Functional annotation analysis of the 1343 genes with overlapping peaks (Table 1; Supplemental Table S7) found approximately one-third enriched the gene ontology *membrane*. Among the genes in this category were nine glutamate receptors, ten solute carriers, the Wnt receptor *Fzd1* and receptors for prolactin, growth hormone, insulin and parathyroid hormone. Within the category GO:0007156∼*homophilic cell adhesion via plasma membrane adhesion molecules* were fourteen cadherin genes and three atypical cadherins. Also among the BMAL1 targets were genes that encoded multiple ephrin receptors and their ligands, prostaglandin E receptors, mitochondrial fission regulators, cell division regulators, multiple steroid hormone receptors and extracellular matrix proteins and proteases (Table 1).

Ingenuity Pathway Analysis (IPA) tools were used to further explore the data set of potential BMAL1 targets represented by overlapping peaks between samples. The most enriched IPA canonical pathway was *Synaptogenesis Signaling Pathway* (*p*=1.21E-10; Supplemental Table S8). Among the 43 genes that enriched this pathway were fourteen cadherins, ephrins and their receptors, glutamate receptors, the very low density lipoprotein receptor (*Vldr*), and alpha synuclein (*Snca*). Another highly enriched canonical pathway was *Axonal Guidance Signaling* (*p-*value=4.67E-6; Supplemental Table S9). Genes that enriched this pathway included two round about guidance receptors (*Robo1, Robo2*) and their ligand (*Slit2*), semaphorins (Sema3A, Sema5A), *Wnt2, Wnt2b,* unc-5 netrin receptors (*Unc5c, Unc5d*) and netrin G (*Ntng1*). IPA identified the glutamate receptor GRIN3A (*p*-value=1.05 E-16; Supplemental Table S10), the CREB transcription factor (*p*-value=2.76 E-14; Supplemental Table S11), and the chorionic gonadotropin complex (CG; *p*-value= 2.28 E-8; Supplemental Table S12) as among the most significant upstream regulators of BMAL1 target genes, with 32, 72 and 45 molecules in the datasets, respectively. The IPA generated regulator network with the highest consistency score indicated the potential BMAL1 targets positively affected processes involved in formation of cellular protrusions, branching of cells, development of sensory organ, sprouting, size of body, and efflux of lipids, while inhibiting organismal death (Supplemental Table 13; Supplemental Figure S2.) A schematic, which represented approximately 15% of the BMAL1 targets, was generated using IPA my pathway tools by combining genes identified as downstream targets of CREB and estradiol (Figure 2; Supplemental Table 14). The schematic illustrates that BMAL1 targets encode proteins across multiple subcellular locations, and the presence of multiple ligand and receptor pairs to include prolactin (*Prl8a2*) and its receptor (*Prlr*), *Slit2* and *Robo*, *Wnt2* and *Fzd1*, glial cell derived neurotrophic factor (*Gdnf*) and its receptor (*Gfra1*) and insulin (*Ins1*) and its receptor (*Insr*).

**Figure 2.**
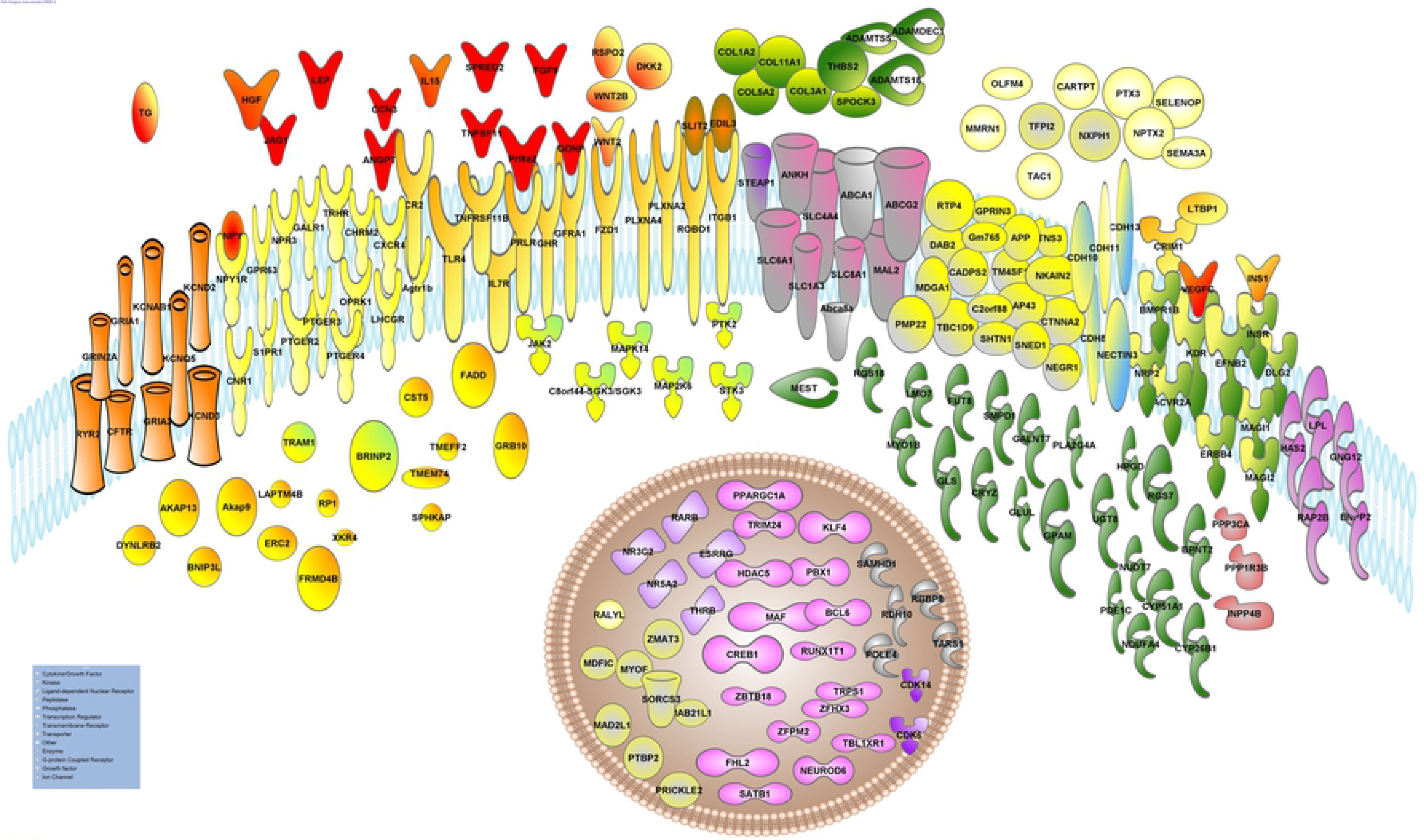
Schematic showing cellular location of BMAL1 targets downstream of CREB1 and estradiol generated using Ingenuity Pathway Analysis tools, with the addition of insulin and Fzd1 genes. Gene names and symbols are given here and can be found with more detail in Supplemental Table S14. ATP-binding cassette subfamily A (Abca1, Abca8a, Abcg2); Activin A receptor (Acvr2a), ADAM metallopeptidase with thrombospondin type 1 (Adamts5, 18) and ADAM like decysin (Adamdec1), angiotensin II receptor (Agtr1b), A kinase anchoring protein (Akap9,13), angiopoietin (Angpt1), inorgancic phosphate transporter (Ankh), amyloid beta precursor protein (APP), bone morphogenetic protein receptor type 1B (Bmpr1b), bisphosphate nucleotidase (Bpnt2) retinoic acid inducible (Brinp2), chromosome 2 open reading frame 88 (C2orf88), serum/glucocorticoid regulated kinase (Sgk3), calcium dependent secretion activator 2 (Cadps2), cellular communication network factor 3 (Ccn3), cadherins (Cdh8, 10,11, 13), cyclin dependent kinase (Cdk6, 14), CF transmembrane conductance regulator (Cftr), cholinergic receptor muscarinic 2 (Chrm2), cannabinoid receptor 1 (Cnr1), collagens (Col1a2, 3a1, 5a2, 11a1), complement C3d receptor 2 (Cr2), cAMP responsive element binding protein 1 (Creb1), cysteine rich transmembrane BMP regulator 1 (Crim1) crystallin zeta (Cryz), cystatin (Cst5), catenin alpha 2 (Ctnna2), chemokine receptor (Cxcr4), cytochrome P450 family (Cyp26b1, 51a1), DAB adaptor protein 2 (Dab2), dickkopf WNT signaling pathway inhibitor 2 (Dkk2), discs large MAGUK scaffold protein 2 (Dlg2), dynein light chain roadblock-type 2 (Dynlrb2), EGF like repeats and discoidin domains 3 (Edil), ephrin B2 (Efnb2), ectonucleotide pyrophosphatase (Ennpp2), erb-b2 receptor tyrosine kinase 4 (Erbb4), ELKS/RAB6-interacting/CAST family member 2 (Erc2), estrogen related receptor gamma (Esrrg), Fas associated via death domain (Fadd), fibroblast growth factor (Fgf9), four and a half LIM domains 2 (Fhl2), FERM domain containing 4B (Frmd4b), fucosyltransferase 8 (Fut8), polypeptide N-acetylgalactosaminyltransferase 7 (Galnt7), galanin receptor (Galr1), growth associated protein 43 (Gap43), glial cell derived neurotrophic factor (Gdnf), growth hormone receptor (Ghr), glutaminase (Gls), glutamate-ammonia ligase (Glul), G protein subunit gamma 12 (Gng12), glycerol-3-phosphate acyltransferase (Gpam), G protein-coupled receptor 63 (Gpr63), GPRIN family member 3 (Gprin3), growth factor receptor bound protein 10 (Grb10), glutamate ionotropic receptors (Gria1, Gria3, Grin2a), hyaluronan synthase 2 (Has2), histone deacetylase 5 (Hdac5), hepatocyte growth factor (Hgf), 15-hydroxyprostaglandin dehydrogenase (Hpgd), interleukin 15 (Il15), interleukin receptor (Il7r), inositol polyphosphate-4-phosphatase type II B (Inpp4b), insulin (Ins1), integrin subunit beta 1 (Itgb1), jagged canonical Notch ligand (Jag1), janus kinase 1 (Jak1), potassium voltage-gated channel (Kncab1, d2, d3, q5), kinase insert domain receptor (Kdr), Kruppel like factor 4 (Klf4), leptin (Lep), luteinizing hormone/choriogonadotropin receptor (Lhcgr), LIM domain 7 (Lmo7), lipoprotein lipase (Lpl), latent transforming growth factor beta binding protein 1 (Ltbp1), mab-21 like 1 (Mab21l1), mitotic arrest deficient 2 like 1 (Mad2l1), MAF bZIP transcription factor (Maf), membrane associated guanylate kinase (Magi1,2), mal, T cell differentiation protein 2 (Mal2), mitogen-activated protein kinase (Map2k6,k14), MyoD family inhibitor domain containing (Mdfic), MAM domain containing glycosylphosphatidylinositol anchor 1 (Mdga1), mesoderm specific transcript (Mest), myosin (Myo1b), myoferlin (Myof), NDUFA4 mitochondrial complex associated (Ndufa4), Nectin cell adhesion molecule (Nectin3), neural growth regulator (Negri), neuronal differentiation 6 (Neurod6), sodium/potassium transporting ATPase interacting (Nkain2), natriuretic peptide receptor 3 (Npr3), neuronal pentraxin 2 (Nptx2) neuropeptide Y (Npy) and receptor (Npr1r), nuclear receptors (Nr3c2-mineralcorticoid receptor; Nr5a2-liver receptor 1, an essential transcriptional regulator of lipid metabolism), neuropilin 2 (Nrp2), nudix hydrolase 7 (Nudt7), neurexophilin 1 (Nxph1), olfactomedin 4 (Ofm4), opioid receptor kappa 1 (Oprk1), PBX homeobox 1 (Pbx1), phosphodiesterase 1C (Pde1c), phospholipase A2 (Pla2g4a), plexin (Plxna2, a4), peripheral myelin protein 22 (Pmp22), PPARG coactivator 1 alpha (Ppargc1a), protein phosphatase 1 regulatory subunit 3B (Ppp1r3b), protein phosphatase 1 catalytic subunit alpha (Ppp3ca), prickle planar cell polarity protein 2 (Prickle2), prolactin (Prl8a2), prolactin receptor (Prlr), polypyrimidine tract binding protein 2 (Prbp2), prostaglandin E receptor (Ptger2,3,4), protein tyrosine kinase 2 (Ptk2), pentraxin (Ptx3), RALY RNA binding protein like (Ralyl), Rap2b, RB binding protein 8 (Rbbp8), retinol dehydrogenase 10 (Rdh10), regulator of G protein signaling (Rgs7, 18), roundabout guidance receptor 1 (Robo1), RP1 axonemal microtubule associated (Rp1), R-spondin 2 (Rspo2), receptor transporter protein 4 (Rtp4), RUNX1 partner transcriptional co-repressor 1 (Runx1t1), ryanodine receptor 2 (Ryr2), sphingosine-1-phosphate receptor 1 (S1pr1), Samhd1, SATB homeobox 1 (Satb1), selenoprotein P (Selnop), semaphorin 3A (Sema3a), shootin 1 (Shtn1), solute carrier (SLC1a3, 4a4, 6a11, 8a1), slit guidance ligand 2 (Slit2), sphingomyelin phosphodiesterase 1 (Smpd1), sushi, nidogen and EGF like domains 1 (Sned1), sortilin related VPS10 domain containing receptor 3 (Sorcs3), SPHK1 interactor, AKAP domain containing (Sphkap), SPARC-osteonectin (Spock3), sprouty related EVH1 domain, Steap1, containing 2 (Spred), serine/threonine kinase 3 (Stk3), tachykinin precursor 1 (Tac1), threonyl-tRNA synthetase 1 (Tars1), Tcb1d9, TBL1X receptor 1 (Tbl1xr1), tissue factor pathway inhibitor 2 (Tfpi2), thyroglobulin (Tg), thyroid hormone beta (Thrb), toll like receptor 4 (Tlr4), transmembrane (Tm4sf1, Tmeff2, Tmem74), TNF (Tnfsf11) and TNF receptor (Tnfrsf11b), tensin 3 (Tns3), translocation associated membrane protein 1 (Tram1), thyroid releasing hormone receptor (Trhr), tripartite motif containing 24 (Trim24), transcriptional repressor GATA binding 1 (Trps1), UDP glycosyltransferase 8 (Ugt8), vascular endothelial growth factor C (Vegfc), Wnt2, Wnt2b, Xkr4, zinc finger (Zbtb18, Zfhx3, Zfpm2, Zmat3).

### Analysis of overlap of protein coding genes closest to ChIP-seq peaks with genes exhibiting circadian rhythms of expression in mammary and liver BMAL1 target genes

A secondary approach to verify BMAL1 targets identified with ChIP-seq, was to query for overlap of targets with genes that exhibited circadian rhythms of expression in mammary tissue or identified as BMAL1 targets in hepatic tissue. The set of genes (Supplemental Tables S2 and S3) used as potential BMAL1 targets was relaxed for this analysis, and although genes were common among two to four ChIP samples the peaks did not necessarily overlap. Genes identified as potential targets in UNDIFF and DIFF samples overlapped with 102 genes that exhibited circadian rhythms of expression in virgin mouse glands[13] and 189 in lactating breast of women[14]. Functional annotation analysis of overlap between mammary circadian rhythm transcriptomes and potential BMAL1 targets categorized thirteen genes in GO:0006974∼*cellular response to DNA damage stimulus* (Supplemental Table S15 and Table 2). Nine genes were clustered in GO:0006629∼*lipid metabolic process* including, *Fdft1, Gpam*, a thromboxane synthase (*Tbxas1*), and several lipases (*Mgll, Plcg2, Plcl2, Pla2g4a*). Other genes overlapping between mammary transcriptomes and potential BMAL1 targets were *Pfkp*, which catalyzes the first step of glycolysis, *Prdx6*, which encodes an antioxidant protein, several homeobox transcription factors (*Sox13, Sox 17, Fox I1*), the mineralcortin receptor (*Nr3c2*), and the transcription factor thyrotroph embryonic factor (*Tef*).

Analysis of overlap of BMAL1 targets in HC11 cells with high confidence BMAL1 targets of protein coding genes in hepatic tissue[19] found 244 in common between liver and mammary epithelial cells. Categories genes enriched included *lipid metabolic process transport* (e.g. transporters for citrate-*Scl13a2*, amino acids-*Slc25a13*, lactate, pyruvate-*Slc16a7*, glucose-*Slc2a2*, and several ABC transporters), *positive regulation of transcription from RNA polymerase II promoter,* and 7 (3%) in *circadian rhythm* (Supplemental Table S16 and Table 2)

### Impact of CRISPR-CAS knockout of BMAL1 on HC11 cell growth, differentiation and metabolic activity

To validate targets and gain an understanding of the functional role that BMAL1 plays in regulation of mammary epithelial cell growth and differentiation, a monoclonal line with the BMAL1 gene knocked out (i.e. deleted; BMAL1-KO) using CRISPR-CAS9 technology was created. PCR analysis prior to monoclonal selection demonstrated the donor cassette was integrated into target sites using both available guide RNAs (gRNA1 and gRNA2; Supplementary Figure S3). Western blot analysis found complete loss of BMAL1 protein expression in several monoclonal lines and supported homozygous knockout of BMAL1 gene was achieved with CRISPR-CAS 9 (Figure 3a). Following Sanger sequencing to confirm genomic integration of green fluorescent protein from the donor cassette (Supplementary Table S17), a monoclonal BMAL1-KO line created using gRNA1 was selected to use for all subsequent studies. Temporal analysis of circadian *Per2* expression indicated 24 hr rhythms of gene expression in wild type HC11 (R^2^= 0.85; *p*-value= 0.00008; amplitude= 1.58) cells, and an attenuation of rhythms in the BMAL1-KO (R^2^= 0.47; *p*-value= 0.04; amplitude= 0.70) line (Figure 3b). Consistent with our previous results[5, 18] expression of BMAL1 protein increased in lactogen DIFF versus UNDIFF HC11 cultures (Figure 3c).

**Figure 3.**
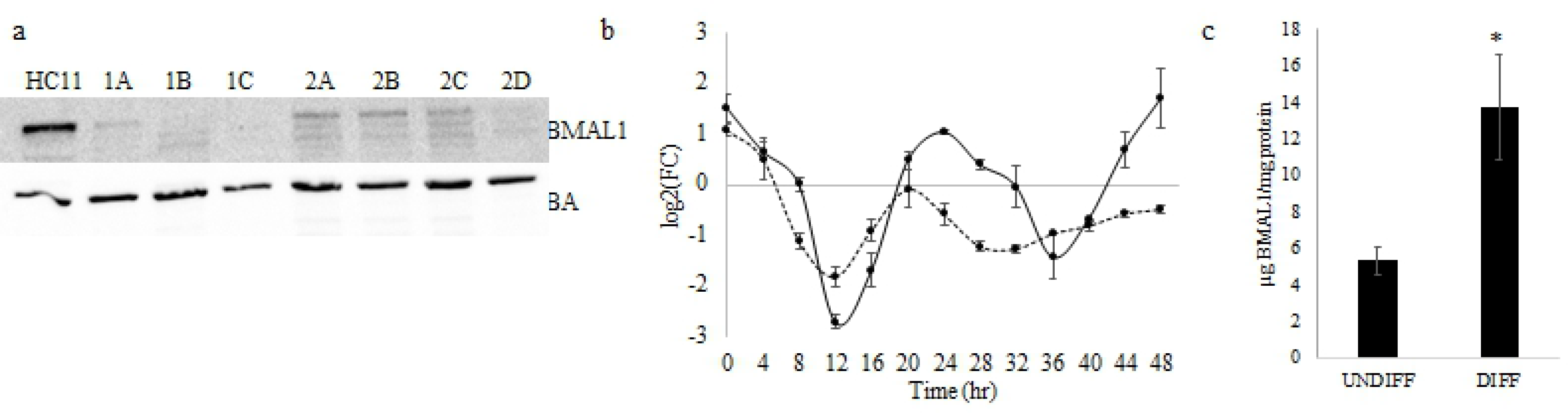
(a)Western blot analysis of BMAL1 protein abundance in HC11 and monoclonal colonies (1A, 1B, 1C, 2A, 2B, 2C, 2D) established from CRISPR-CAS transfected with guide RNA (gRNA) targeting BMAL1 gene post monoclonal selection. BMAL1 is absent in colonies 1C and 2D and decreased in the remaining colonies compared to WT HC11. Monoclonal 1C culture was used in all subsequent experiments, and referred to as BMAL1-KO. Data are representative of two western blots. (b) Temporal analysis of *Per2* expression in WT HC11 (solid line) and BMAL1-KO (dashed line) cultures. For this experiment cells were grown to confluence in growth media. Media was changed to lactogen media for 2 hr to synchronize clocks. At completion of 2 hr lactogen treatment (time 0 hr), cells were rinsed with PBS and cultured in growth media for remainder of the experiment. Cells were collected for isolation of total RNA every 4 hr over a 48 hr period beginning at 0 hr. *Per2* was measured with q-PCR, and levels were expressed relative to mean levels across all time points of HC11 culture. Cosinor analysis found mesor (−0.12 and −0.68), amplitude (1.58 and 0.70), acrophase (0.78 and −0.56), R^2^ (0.85 and 0.47) and *p*-value (8.05 E-5 and 0.04) of fit to a 24 hr rhythm were calculated, respectively, for HC11 and BMAL1-KO lines. Data represent n=3 wells/line and 2 experimental replicates. (c) BMAL1 protein abundance in UNDIFF and DIFF cultures measured using ELISA. Data are expressed as mean µg of BMAL1/mg protein ± standard deviation of three samples per treatment; * indicates difference at *p*<0.05. Data represent n=3 protein isolates/line/state of differentiation.

Eight-day growth curve analysis showed that although doubling time between wild-type HC11 and BMAL1-KO lines was not different (*p*>0.05), with HC11 at 21.9 ± 3.8 hr and BMAL1-KO at 26.9 ± 3.8 hr, the BMAL1-KO line reached stationary phase at a significantly lower cell density than wild-type HC11 cells (*p*<0.05; Figure 4a). The MTT assay found a similar pattern as the eight-day growth curve (Figure 4b). Analysis of images captured on days 2 and 6 of culture (Figure 4d) found less cells and lower (*p*<0.05) intensity of MTT staining per unit area of cells in BMAL1-KO cultures (Figure 4c), indicating that BMAL1-KO cultures had less cells and lower metabolic activity per cell than wild-type HC11 cultures. FACS analysis found no difference between HC11 and BMAL1-KO in the proportion of cells in G1/G0 and S/G2/M phase of the cell cycle across the eight days (Figure 4e). FACS screening for cells with less than 2N, an indicator of dead or dying cells, found a higher percent in BMAL1-KO cultures on all eight days (*p*<0.05; Figure 4f). Despite no difference in the percent of cells in S/G2/M phases, the expression level of *Ccnd1*, which regulates the transition from G1 to S phase of the cell cycle, was greater (*p*<0.05) in BMAL1-KO line (Figure 4g).

**Figure 4.**
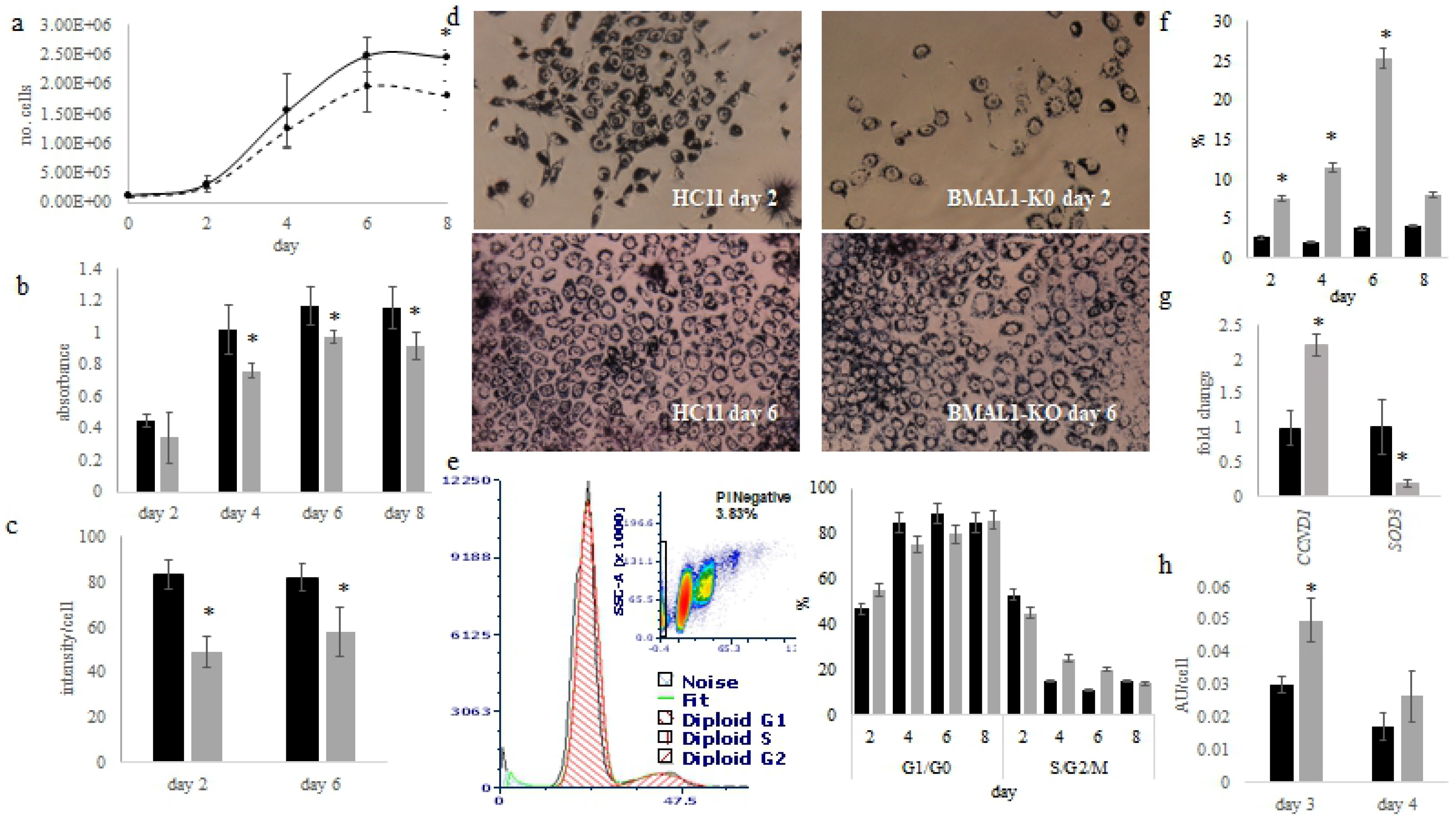
Effect of BMAL1 CRISPR-CAS knockout (BMAL1-KO) on growth and metabolic activity in culture. (a) Eight day growth curve analysis was performed by plating 100,000 cells/ well on day 0 in 6 well dish of wild type HC11(black line) and BMAL1-KO (gray line), two wells/treatment were collected and counted every two days; values are mean ± standard deviation across five experiments. Two way ANOVA found that line and day significantly affected (*p*<0.05) number of cells. Data represent five experimental replicates, with n=2 wells/line per experiment. (b) MTT assay was performed by plating HC11 (black bars) and BMAL1-KO (gray bars) cells at 10,000 cells/well in a 96-well plate; on days 2, 4, 6 and 8 of culture, MTT assay was performed. ANOVA found line and day significantly affected NADH levels; values are mean ± standard deviation; * indicates difference between lines at *p*<0.05 across 3 replicate experiments. Data represent three replicate experiments collected from n=3 wells/line/day. (c) The intensity of MTT staining per cell across three images on each day in HC11 (black) and BMAL1-KO (gray) cultures was quantified. Values are mean intensity per cell ± standard deviation. A significant difference at *p*<0.05 is indicated by *. (d) Images of cells were captured following staining with the MTT assay on day 2 and 6 of culture. (e) Cells were collected from 100 mm dishes (plating density was 100,000 cells/ml) for fluorescence activated cell sorting (FACS) to determine percent of cells in G1/G0 and S/G2/M phases following labeling with propidium iodide across 8 days of culture. Values are mean across 5 experiments, with ANOVA analysis finding that day had an effect (*p*<0.05) on proportion of cells in phases, but there was no difference between HC11 (black) and BMAL1-KO (gray). (f) FACS analysis for dead or dying cells (cells or events with <2N) in HC11 (black) and BMAL1-KO (gray). Values are percent of total events ± standard error; * indicates difference at *p*<0.05. (g) q-PCR analysis of *Ccnd1* and *Sod3* expression in undifferentiated (UNDIFF) cultures of HC11 (black) and BMAL1-KO (gray) cells. Values are mean across triplicate samples and two experimental replicates, normalized to express fold-change relative to mean of HC11 ± standard deviation using delta-delta cycle threshold method; Student t-test analysis * indicates difference between lines at *p*<0.05. (h) Reactive oxygen species (ROS) assay of HC11 and BMAL1 cells on day 3 and 4 of culture. Two-way ANOVA found day and line affected (*p*<0.05) ROS levels; * indicates difference between lines at *p*<0.05 across 3 replicate experiments n=3 wells/line/day; values are mean of arbitrary units (AU) ± standard deviation.

Others reported that transgenic mice with BMAL1 gene knocked out (*Bmal1^−/−^*) had reduced lifespans and displayed symptoms of premature aging associated with increased levels of reactive oxygen species in some tissues[20]. To determine if this was the potential cause of cell loss in BMAL1-KO line, reactive oxygen species (ROS) were measured and found to be significantly higher in BMAL1-KO cultures on days measured (Figure 4h). Q-PCR analysis of the antioxidant *Sod3*, which was identified as a potential target of BMAL1 in HC11 cells, revealed that mRNA levels were significantly lower in BMAL1-KO versus wild-type HC11 cultures (Figure 4g).

In the DIFF2 sample the serotonin transporter-SERT (*Slc6a4*) and tryptophan hydroxylase 1 (*Tph1*), which encodes the protein that catalyzes the first and rate-limiting step in the biosynthesis of serotonin, were identified as potential targets. Lactogen induced differentiation of HC11 cells significantly increased mRNA levels of *Tph1* (Figure 5a) and *Sert* (Figure 5b), whereas levels were significantly lower in the BMAL1-KO line. Q-PCR analysis using primers that targeted two sites in the *Sert/Slc6a4* promoter region that contained E-box sequences beginning at −42 and −1282 nucleotide bases upstream of the transcriptional start site, found levels 3.5-fold and 2-fold, respectively, higher in UNDIFF BMAL1 ChIP samples than mock-IP samples (Figure 5c). Although only one DIFF sample had a peak in the promoter region of the *Sert/Slc6a4* gene (Supplemental Figure S4f).

**Figure 5.**
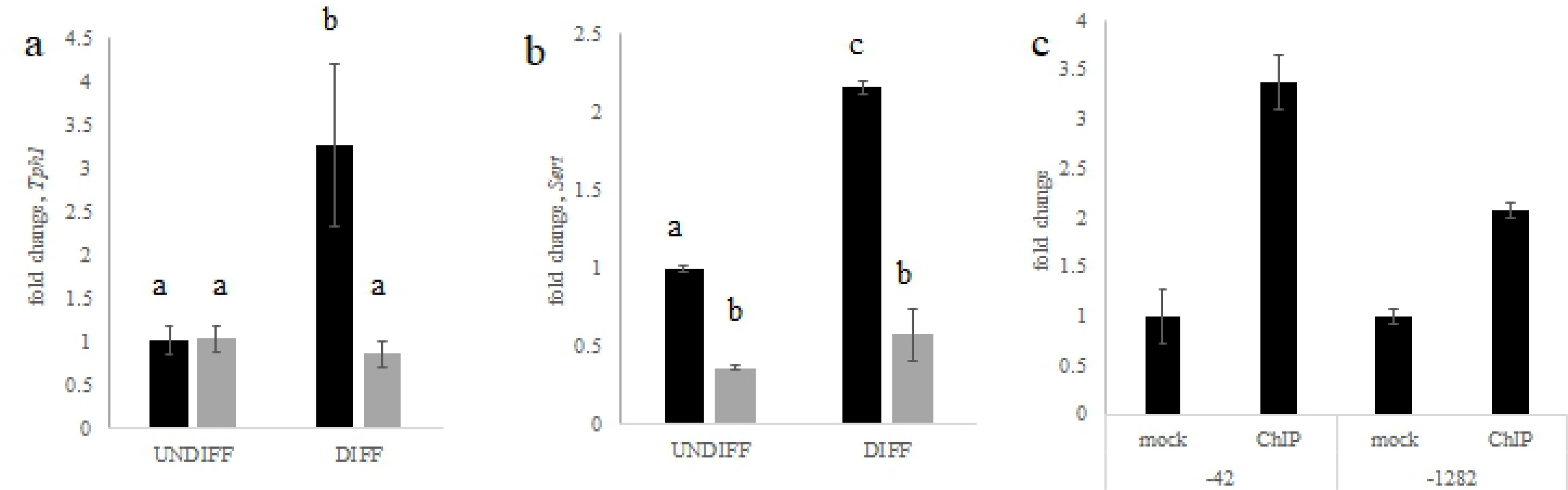
Quantitative-PCR (q-PCR) analysis of (a) *Tph1* and (b) *Slc6a4 (Sert)* in undifferentiated (UNDIFF) and lactogen differentiated (DIFF) HC11 (black) and BMAL1-KO (gray) cultures. Values are mean across triplicate samples and two experimental replicates, normalized to express fold-change relative to mean of HC11 ± standard deviation using delta-delta cycle threshold method; ANOVA and post-hoc Tukey test analysis indicated with differing letter reflecting difference at *p*<0.05. (c) qPCR analysis of *Slc6a4 (Sert)* promoter region using primers that targeted two sites that contained E-box sequences beginning at −42 and −1282 nucleotide bases upstream of the transcriptional start site in undifferentiated HC11 cultures; values are mean across four samples, normalized to express fold-change relative to mean of mock ± standard deviation. A 2-fold difference was considered a positive ChIP.

*Ppara*, which regulates lipid metabolism and glucose homeostasis, was also identified as a potential target in both DIFF samples (Supplemental Figure S4a and S4b) as well as in the liver[19]. Q-PCR analysis of *Ppara* expression levels found mRNA depressed in UNDIFF and DIFF BMAL1-KO cultures relative to wild-type HC11 (Figure 6a). Fatty acid synthase (*Fasn*) was identified as a potential target of BMAL1 in both DIFF samples (Supplemental Figure S4e). However, q-PCR analysis across experimental replicates showed variable results (Supplemental Figure S5a-c). Moreover, temporal of *Fasn* mRNA expression across a 48 hr period indicated lack of a fit (*p*>0.05) to a 24 hr rhythm (Supplemental Figure S5c). Therefore, it was concluded that *Fasn* mRNA levels were not different between BMAL1-KO and HC11 lines.

**Figure 6.**
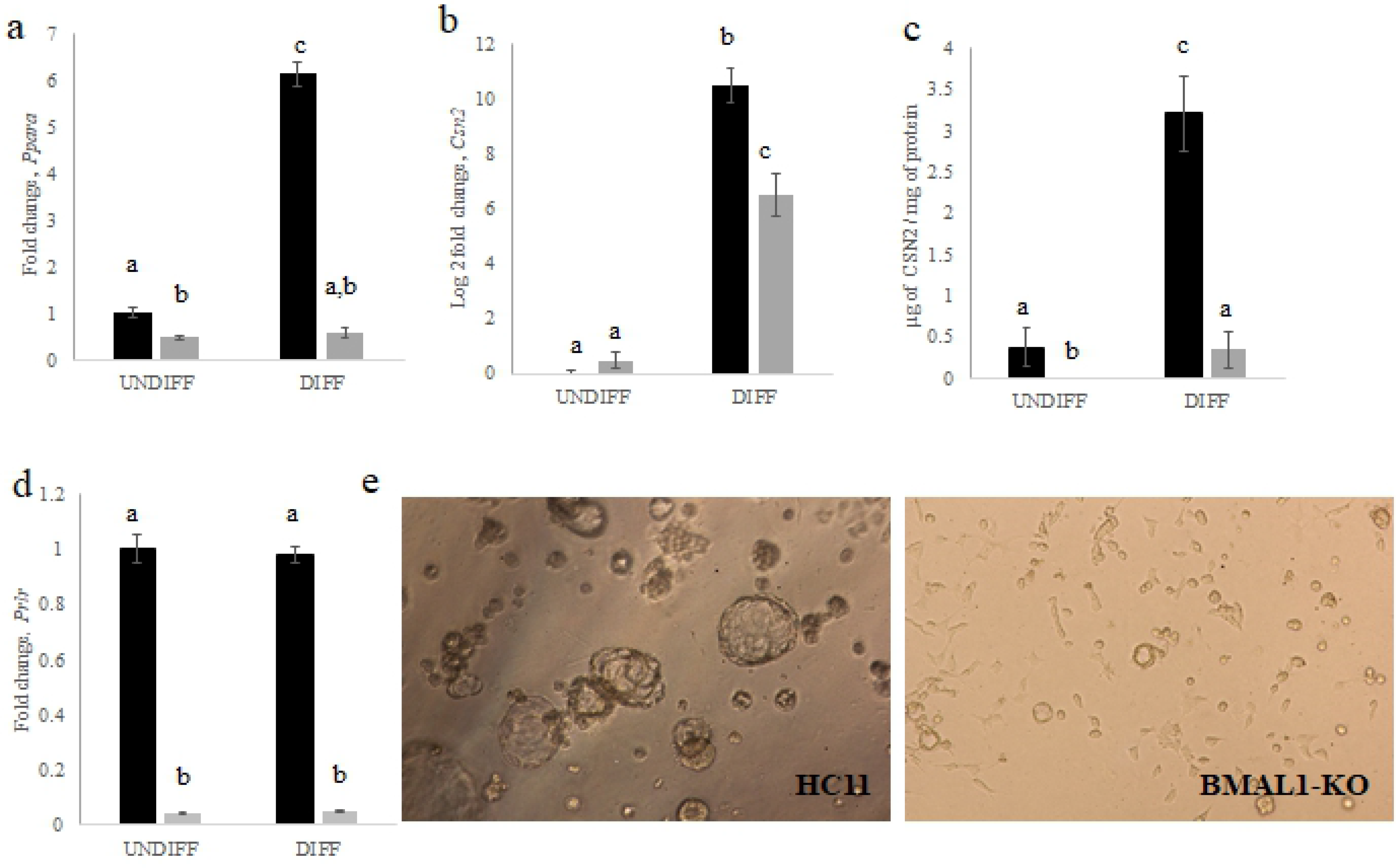
Quantitative-PCR (q-PCR) analysis of (a) *Ppara* and (b) *Csn2* in undifferentiated (UNDIFF) and lactogen differentiated (DIFF) HC11 and BMAL1-KO cultures. Values are mean across triplicate samples and two replicate experiments, normalized to express fold-change relative to mean of HC11 ± standard deviation using delta-delta cycle threshold method. Note the y-axis in *Ppara* is fold change, whereas y-axis for *Csn2* is log base 2 of fold change due to large induction. ANOVA and post-hoc Tukey test analysis findings is indicated by differing letter reflecting difference at p<0.05. (c) ELISA quantification of CSN2 protein in UNDIFF and DIFF HC11 (black) and BMAL1-KO (gray) cultures. Values are mean concentration ± standard deviation across triplicate samples. ANOVA and post-hoc Tukey test analysis findings is indicated by differing letter reflecting difference at p<0.05. (d) q-PCR analysis of *Prlr* in UNDIFF and lactogen DIFF HC11 (black) and BMAL1-KO (gray) cultures. Values are mean across triplicate samples, normalized to express fold-change relative to mean of HC11 ± standard deviation using delta-delta cycle threshold method. Note the y-axis is fold change. ANOVA and post-hoc Tukey test analysis findings is indicated by differing letter reflecting difference at p<0.05. (e) Images of two and a half dimensional drip gel cultures of HC11 and BMAL-KO cells taken with phase-contrast microscopy after 7 days of incubation in lactogen media. Cells were plated at 13,000 cells/well.

Although *Csn2*, which encodes the milk protein beta-casein, was only identified as a potential target in one of the differentiated samples (DIFF2), levels of mRNA (Figure 6b) and protein (Figure 6c) were lower in BMAL1-KO versus wild-type HC11 lines. Prolactin regulates *Csn2* expression and the prolactin receptor (*Prlr;* Supplemental Figure S4c and S4d) was identified as a BMAL1 target. *Prlr* mRNA expression was reduced in UNDIFF and DIFF BMAL1-KO cultures (Figure 6d). Lower *Prlr* levels would affect multiple pathways that stimulate mammary epithelial cell differentiation. The ability of mammary epithelial cells to differentiate can be evaluated by acini formation in Matrigel, a laminin rich extracellular matrix extract.[21] After seven days in culture, wild-type HC11 cells formed many relatively large acini, whereas BMAL1-KO cultures primarily failed to do so, and acini that were formed were much smaller in size (Figure 6e).

## DISCUSSION

Genes identified in undifferentiated and lactogen differentiated mammary epithelial cells as potential transcriptional targets of BMAL1 support the hypothesis that the circadian clock in the mammary gland regulates mammary development and lactation, and that transcriptional targets of BMAL1 change as mammary epithelial cells differentiate. The greater number of transcriptional targets in differentiated HC11 cultures were likely due, in part, to the effects of lactogenic hormones on chromatin access. Higher doses of glucocorticoid treatment are associated with pioneering activities of glucocorticoid receptors. Binding of activated glucocorticoid receptors to chromatin increases the accessibility of regions of the genome previously inaccessible to other transcription factors.[22, 23]. Prolactin (*Prl8a2*) and the prolactin receptor (*Prlr*) were identified as BMAL1 targets. Prolactin regulates *Bmal1* expression in mammary epithelial cells [5], and mammary epithelial cells synthesize and secrete prolactin [24]. Thus, in differentiated mammary epithelial cells BMAL1 may play a role in the iterative induction of its own activity through positive loops with prolactin signaling. Upon prolactin binding to its receptor JAK2 is activated. *Jak2* was also identified as a BMAL1 target. Activated JAK2 phosphorylates and activates STAT5 transcription factors. Activated STAT5 transcription factors were associated with pioneer-like effects on chromatin in mammary epithelial cells [25]. Thus, the higher number of BMAL1 targets in the lactogen differentiated cultures may have been due to dexamethasone and prolactin treatments increasing the accessibility of BMAL1 to chromosomal regions. Moreover, the positive loop between BMAL1 and prolactin signaling molecules may potentially explain the changes in stoichiometric relationships between positive and negative elements of the mammary epithelial clock transcription-translation feedback loop observed following lactogen induced cellular differentiation [5].

The prediction of CREB as an upstream regulator of BMAL1 targets is consistent with CREB’s role as a coactivator of BMAL1:CLOCK transcription factor activity [26, 27]. Genes identified as CREB targets encompassed cell adhesion molecules (cadherins, catenin and nectin genes), leptin, lipoprotein lipase, and estrogen receptor gamma. The prediction of the glutamate-regulated ion channel GRIN3A as the most significant upstream regulator of BMAL1 target genes, suggest that, similar to the role that glutamate places in regulation of the master clock in the SCN [28], it potentially entrains circadian rhythms in mammary epithelial cells. Moreover, the identification of chorionic gonadotropin complex as an upstream regulator of BMAL1 target genes may reflect a potential role of the mammary epithelial clock in integration and coordination of the timing of endocrine-paracrine signal-receptivity-response. Among the targets downstream of this complex were receptors for mineralcorticoid (*Nr3c2*), prolactin (*Prlr*), prostaglandin E (*Ptger2*), natriuretic peptide (*Npr3*), luteinizing hormone/choriogonadotropin (*Lhcgr*), bone morphogenetic protein (*Bmpr1b*). In addition, there were receptors for Wnt ligands (*Fzd1*), ephrins (*Epha3* and *Epha7)*, an integrin (*Itgb1*), and the orphan nuclear receptor *Nr5a2*, which acts as a metabolic sensor that regulates expression of genes involved in bile acid synthesis, cholesterol homeostasis and triglyceride synthesis.

Among the BMAL1 targets identified were multiple components of the Wnt signaling pathway (*Wnt2, Wnt2b, Fzd1, Rspo2, Ctnna2*) and *Cdk14*, which acts as a cell-cycle regulator of Wnt signaling pathway. The Wnt signaling pathway influences mammary stem cells to elicit fate specification and patterning during embryonic and postnatal development of the gland[29]. Regulation of the Wnt/β-catenin pathway by the circadian clock has been demonstrated in epidermal stem cells [30]. Alterations in expression level of genes in the Wnt pathway may potentially be the cause of compromised mammary epithelial stem cell function observed in *Clock-Δ19* mice[13], as well as the premature aging evident in *Bmal1-/-* mice[31]. The nuclear hormone receptors for glucocorticoids, retinoids, and estrogens also regulate adult stem cell fate [31]. Retinoic acid receptor beta (*Rarb*), the orphan receptor estrogen-related receptor gamma (*Esrrg*), which interferes estrogen receptor signaling, and the mineralcorticoid receptor (*Nr3c2*), which binds glucocorticoids, were found among the potential targets of BMAL1 in mammary epithelial cells. Additional evidence that the clock in mammary epithelial cells may regulate cell fate and developmental patterning was the network generated by IPA software that indicated BMAL1 targets affected branching of cells, development of organs, and body size, while inhibiting cell death. Moreover, numerous BMAL1 targets were involved in stem cell maintenance to include *Sox1* and *Sox6* [32] and *Pard3* and *Pard3b* genes. *Pard3* is necessary for mammary gland morphogenesis, and *Pard3b* is expressed by multipotent stem cells in the terminal end buds of murine mammary glands, with studies finding ablation of *Pard3b* resulted in stem cell loss [32].

Also among the BMAL1 targets were a multitude of ion transporters, solute carriers, glutamate transporters and ATP-binding cassette transporters. Synthesis and secretion of milk is dependent on membrane transport systems that move ions and substrates into and out of epithelial cells [33, 34]. Movement of ions by ion transporters creates potential differences that enable electrical signaling which regulates cell number, shape, differentiation, and morphogenesis [35]. Glutamine is the primary nitrogen donor for proliferating cells, and alterations in intracellular glutamine abundance was associated with quiescent and proliferative states of mammary epithelial cells [36]. Glutamine also contributes carbons to biosynthetic reactions, and thus mammary clock regulation of glutamate transport may be a means by which the timing of proliferation and metabolic activities of epithelial cells are coordinated. Of the ten ATP binding cassette genes identified as BMAL1 targets seven were the ABCA subfamily type, which function to mediate efflux of cholesterol (*Abca1*) and other lipids from cells. Others included *Abcc12,* which mediates toxin efflux, the xenobiotic transporter *Abcg2,* and *Abcb5,* which may also function in the efflux of drugs and toxins. Thus, enrichment of pathways and categories related to ion channels, synapse and substrate transports point to the potential role of the mammary clock in affecting the program of mammary development as well as regulating substrate availability and detoxification during lactation.

Functional studies with the BMAL1-KO line found that deletion of *Bmal1* did not affect proliferation rate of cells, but rather resulted in greater rates of cell death and lower metabolic activity. Although, *Ccnd1* was identified as a potential BMAL1 target, deletion of *Bmal1* gene resulted in significantly higher expression of *Ccnd1* in BMAL1-KO cultures. Similarly, when abundance of CLOCK protein was decreased in HC11 cells with shRNA, *Ccnd1* expression was higher and proliferation rate increased[18]. In our previous study, lower CLOCK abundance appeared to result in the loss of gating of entry and progression through the cell cycle. Elevated *Ccnd1* likely reflected this phenomena in culture, and thus loss of gating may also explain elevated *Ccnd1* levels in BMAL1-KO cells. Gating the time cells enter the cell cycle by clocks enables the sequestering of processes incompatible with DNA synthesis[37]. Loss of gating time, coupled with increased levels of reactive oxygen species, related to lower *Sod3* expression, in BMAL1-KO cells may have resulted in genotoxic stress, and cell death. There were 31 BMAL1 targets that encoded proteins involved in response to DNA damage, to include *Cdkn2aip, Nsmce2, Rad23b, Hus1, Ubr5, Ube2e2, Ube2w.* Thus, death of BMAL1-KO cells was also potentially due to decreased ability to repair DNA damaged by reactive oxygen species. Moreover, *Ubr5, Ube2e2, Ube2w* are involved in the ubiquitination pathway. The ubiquitination pathway targets proteins for degradation and plays a central role in maintaining proteostasis, the cellular balance of protein synthesis and degradation. The aging process is related to changes in proteostatic equilibrium and build-up of misfolded and damaged protein aggregates[38, 39], and thus the observation that Bmal1 knockout mice (*Bmal1* −/-) exhibited advanced aging[20], may be due to alterations in these genes and the ubiquitination pathway.

Involvement of BMAL1 targets in proteostasis supports that the mammary epithelial clock regulates cellular-tissue homeostasis. During lactation, the serotonergic system plays a central role in regulating mammary epithelial cell homeostasis [40]. Multiple components of the serotonergic system were identified as potential BMAL1 targets in several of the samples. TPH1 and SERT/SLC6A4 are central to serotonergic regulation of mammary epithelial homeostasis through regulation of synthesis and degradation of serotonin, respectively. Our previous analysis found multiple E-box sequences in the promoter region *Sert/Slc6A4* and *Tph1* genes, and that SERT exhibited circadian rhythms of expression in HC11 cells and lactating sheep mammary[41]. ChIP-qPCR analysis of regions encompassing E-box sequences in the promoter regions of SERT confirmed BMAL1 binding to these sites in HC11 cells. Levels of *Sert* and *Tph1* were also significantly depressed in BMAL1-KO cultures relative to wild-type HC11. The circadian system functions in part to maintain metabolic homeostasis[42], and thus a role for the mammary clock in regulating factors that control epithelial homeostasis during lactation is consistent with this function.

*Lipid metabolic process* was enriched with BMAL1 target genes, and these overlapped with genes that showed circadian rhythms of expression in mammary and those that were confirmed targets of BMAL1 in hepatic tissue. Notable among them were *Fdft1, Gpam, Vldlr, Plin5* and *Mgll,* all of which increase significantly upon secretory activation of the mammary gland[43, 44], a process marked by the onset of milk fat synthesis and secretion. FDFT1 (farnesyl-diphosphate farnesyltransferase 1) regulates cholesterol synthesis, GPAM catalyzes the synthesis of glycerolipids, VLDLR and MGLL function in the uptake and hydrolysis of triglycerides, and PLIN5 mediates lipid droplet formation. Together supporting that the mammary epithelial clock plays a central role in the regulation of milk fat synthesis during lactation.

Also central to lipid metabolism is PPARA (peroxisome proliferator-activated receptor-alpha). As a nuclear receptor it senses hormonal and nutrient status of the cell and functions to stimulate uptake, utilization, and catabolism of fatty acids[45]. Q-PCR analysis of *Ppara* expression showed that levels were significantly depressed in UNDIFF and DIFF cultures of BMAL1-KO cells versus wild-type HC11, and failed to increase following four days of lactogen treatment. These findings are consistent with the knowledge that PPARA is a well characterized target of the BMAL1:CLOCK transcription factor and believed to be central to coordinating cellular energy metabolism with molecular clocks[45]. The interaction of the mammary clock and PPARA likely explain changes observed in milk composition that occurred with night restricted feeding in dairy cattle[46]. The transport of nutrients also appears to be under the control of the mammary epithelial clock as multiple transporters showed overlap with high confidence BMAL1 targets in hepatic tissue to include citrate, amino acid, lactate, pyruvate, and glucose transporters, and thus potentially also affected circadian rhythms of milk component synthesis.

Fatty-acid synthase (FASN) is the enzyme that catalyzes the first committed step in fatty-acid biosynthesis, and products of FASN mediated synthesis serve as PPARA ligands[47, 48]. Despite FASN being identified as a BMAL1 target in HC11 cells, there was lack of a significant effect of deletion of BMAL1 on *Fasn* expression. This finding maybe due to the integrated and reciprocal regulation of FASN products and PPARA activation[47–50].

BMAL1 expression is significantly increased during the transition from pregnancy to lactation in mouse mammary glands, and this increase is modelled in lactogen treatment of HC11 cells (results and [5]). The increase in BMAL1 is likely a direct response to increased prolactin and prolactin induced signaling in lactogen treated cells and at the onset of lactation, as discussed above. The prolactin receptor (*Prlr*) was identified as a BMAL1 transcriptional target in HC11 cells, and levels of mRNA were significantly depressed in undifferentiated and lactogen differentiated BMAL1-KO cultures. Lower levels of *Prlr* would affect all pathways regulated by prolactin, which is a key hormone in induction of mammary epithelial cell growth, differentiation and milk component synthesis[51]. The lower expression of the milk protein beta-casein (*Csn2*) in DIFF BMAL1-KO cultures, and the reduced ability of this line to form acini in culture were likely due, at least in part, to the lower level of prolactin receptors.

*Cell adhesion, cell junction* and *proteolysis* were categories highly enriched with BMAL1 targets identified in HC11 cells. Cell-cell and cell-extracellular matrix (ECM) interactions play a central role in mammary morphogenesis and regulation of milk synthesis[41]. BMAL1 targets within cell adhesion included fourteen cadherin molecules and three atypical cadherins. Cadherins make-up the extracellular and transmembrane component of adherens junctions and function as cell-cell adhesion receptors that mediate interactions between adjacent cells. The cytoplasmic component of adherens junctions is a multiprotein complex that links adherens junctions with the actin cytoskeleton[52]. The actin cytoskeleton is hypothesized to function to relay SCN-driven timing cues to gene expression in peripheral tissues[53]. Thus, it is not surprising that the mammary clock would control cellular components that connect to factors that transmit temporal information within cells. The dramatic decreased ability of BMAL1-KO cells to form acini in culture likely reflect loss of activity in cell-cell and cell-ECM adhesion molecules that are critical to the morphogenesis and differentiation of mammary epithelial cells. Cell-cell and cell-ECM junctional complexes sense environmental changes and function to maintain mammary epithelial homeostasis[54]. Changes in mechanical tension affect circadian clocks in mammary glands of virgin mice[13]. Adhesion molecules and proteolytic enzymes can affect the environment of the cells, and it has been hypothesized they mediate circadian timing in the central nervous as they can sense rapid signaling events and transmit-transfer signals into broader changes in cell activity with the daily changes[55].

There are several limitations to our studies. When the frequency of peak location of BMAL1 targets found in our study were compared with findings of others, the rate of intergenic sites was higher in our data set and the frequency of peaks found in promoter regions was lower [56]. The frequency of regions of peaks identified were consistent across the four samples analyzed in our study. Therefore, the difference may be tissue or cell type specific, as our studies were conducted using a mammary epithelial cell line, whereas the other group conducted studies using liver tissue from mice. Additionally, to cast a wider net for identification and functional annotation of BMAL1 target genes, the stringency for peak calling was relaxed from the default settings of HOMER, and in doing so the number of false-positives likely increased. To increase the confidence in BMAL1-targets, genes with overlapping peaks were highlighted in the manuscript. When genes identified in our study were compared with datasets of genes that showed circadian rhythms of gene expression in mammary tissue or identified as BMAL1 targets in hepatic tissue, overlapping between datasets was only realized when criteria were further relaxed to common genes identified, but peaks did not necessarily overlap between samples. Similarly, only when this more relaxed selection criterion was used, were BMAL1 (ARNTL) and CLOCK identified as significant upstream regulators predicted by IPA.

Another limitation to the studies described here, is that circadian rhythms of gene expression and BMAL1 binding activity were not measured. Although robust circadian rhythms of core clock genes are evident in UNDIFF HC11 cells, constant exposure to lactogenic hormones, which is required to differentiate cells, results in loss of circadian rhythms of expression of multiple core clock genes [4, 5]. Loss of circadian rhythms of core clock genes’ expression also occurred in mice in early lactation and resulted in relatively constant levels of BMAL1:CLOCK transcription factor. This dynamic is likely due to the hormonal milieu of the physiological state[5]. Lack of capturing a circadian rhythm of BMAL1 binding may have led to our findings that expected targets such as *Per1* were not identified across all four samples (Supplemental Figure S4g). Additionally, data were completely generated using a mammary cell line, and need to be confirmed and studied *in vivo*.

Caution must also be used in interpreting comparisons between HC11 and BMAL1-KO lines. An appropriate negative control was not used for studies as off target effects were observed when the scrambled sequence was transfected into HC11 cells. Off target effects are common with CRISPR-CAS [57], and so findings presented may not be specific to loss of BMAL1 function. Finally, the assay used to measure reactive oxygen species level may not truly be reflective of its content [58], and thus caution is warranted in interpreting these data. Regardless, data presented in the manuscript will serve as a good resource to inform future studies aimed at understanding the role of the mammary clock in mammary morphogenesis and lactation as well as provide insight into the link between circadian disruption and breast cancer and poorer milk production.

Overall, ChIP-seq analysis revealed potential BMAL1 transcriptional targets in mammary epithelial cells. Knowledge of these targets provides insight into the role of circadian clocks in undifferentiated and differentiated states of mammary epithelial cells. BMAL1 transcriptional targets play central roles in pathways that affect stem cell maintenance, cellular detoxification and proteostasis, substrate transport and milk fat synthesis. There was also evidence for the coordination of endocrine-paracrine signals by the mammary epithelial clock, as BMAL1 targets included both the ligand and corresponding receptor as in the case of prolactin, insulin and Wnts with Fzd receptor. In vivo studies are now needed to understand the roles of mammary epithelial clocks in stage specific development of the gland in vivo, with interaction of stromal tissue and physiological context maintained.

## MATERIALS AND METHODS

### Routine culture and differentiation of cell cultures

HC-11 cells were routinely cultured in complete growth medium: RPMI 1640 (50-020-PC, Mediatech Inc.) supplemented with 2 g/L sodium bicarbonate (S5761-500G, Sigma Life Science), 100 U/mL penicillin, 100 µg/mL streptomycin (15140-122, ThermoFisher Scientific), 10% heat inactivated calf serum (26170-043, Gibco), 5 µg/mL insulin ([Ins] IO516-5mL, Sigma-Aldrich), and 10 ng/mL epidermal growth factor ([EGF] E4127, Sigma-Aldrich) in 5% CO_2_ at 37°C. Cells were passaged 1:3 when they reached 80% confluence by harvesting cells with 0.25% trypsin EDTA (Gibco).

To induce differentiation, cells were plated at 100,000 cells/mL and grown to confluence. At confluence cells were washed 2-times with PBS, and incubated for 48 hr in RPMI media supplemented with 10% serum and Ins (no EGF in media). Undifferentiated cultures (UNDIFF) were harvested for downstream analysis at this stage of culture. For differentiated cultures (DIFF), media was changed to RPMI supplemented with 10% calf serum, dexamethasone (0.1 µM), insulin (5 µg/mL), and prolactin (5 µg/mL; ovine prolactin: L6520-250IU, Sigma-Aldrich), and incubated for 96 hr, with media change every 2-days. After 96 hr in lactogen cocktail, differentiated cultures were collected.

For growth curve analysis, 100,000 cells were plated using complete growth media in 6-well dishes, and cells from 2-dishes per time point were harvested using trypsin EDTA and counted on days 2, 4 and 8 using a BioRad TC10 Automated cell counter (BioRad Laboratories, Inc). Growth curves were conducted 5-times, and results were presented as mean number of cells each day ± standard deviation. Doubling time was calculated using the tool available here[59].

To measure circadian rhythms of gene expression, cells were grown to 80% confluence in 6 well plates. Media was changed to lactogen media, and cells were incubated for 2 hr at 37 °C to synchronize clocks[5]. After completion of 2 hr lactogen treatment [designated as circadian time 0 (CT0)], three wells of cells of each line were rinsed with PBS and RLT buffer from the QIAGEN RNeasy kit was added to cultures and lysates were collected and stored at −80 °C for subsequent isolation of total RNA. Cells in remaining wells were washed three-times with PBS, and complete growth media was added. Beginning from CT0 and every 4 hr over the next 48 hr, cells were rinsed with PBS, RLT buffer was added, and lysates collected and stored at −80 °C for subsequent isolation of total RNA.

### Two-and a half dimensional drip cultures and immunofluorescence

RPMI growth media supplemented with 5% serum, 5 μg/mL prolactin, 5 μg/mL insulin, and 0.375 ng/μL hydrocortisone (cat. no. 65966, BD Biosciences) was used for 2.5-dimensional drip cultures to study acini formation[60]. Briefly, cold Matrigel (Growth Factor Reduced: 356231, Corning Life Sciences; 50 μL of gel/cm3) was used to coat the bottom of each well of 4-well chamber plates (Lab-Tek, Nunc). The 4-well chamber plate was placed in 5% CO_2_ at 37°C for 30 minutes to allow the Matrigel to solidify, and then cells were plated at a density of 13,000 cells/cm^2^. Cells were allowed to attach Matrigel for 15 min and then media with 5% Matrigel was added drop by drop to cover cultures. Media was changed every two days. On day seven of cultures, images of cultures were captured at 200 X magnification (20 X objective) with phase contrast on a Zeiss AxioVert (Oberkochen, Germany) inverted microscope using a Nikon camera.

### Creation and screening of CRISPR-CAS BMAL1 knockout HC11 lines

ORIGENE’s ARNTL Mouse Gene Knockout Kit (CRISPR; CAT#: KN301604; Rockville, MD US) was used to knockout (remove) the BMAL1 gene from HC11 cells following manufacturer’s protocol. The kit contained: KN301604G1, *Arntl* gRNA vector 1 (gRNA1) in pCas Guide CRISPR vector with the target sequence: GAACCGGAGAGTAGGTCGGT; KN301604G2, Arntl gRNA vector 2 (gRNA2) in pCas Guide CRISPR vector with the target sequence: CATGAAGTCGCTGATGGTTG; KN301604-D, donor DNA containing left and right homologous arms and green fluorescent protein-puromycin resistance gene functional cassette; and GE100003 scramble sequence in pCas-Guide vector. HC11 cultures in 6-well dishes were plated overnight and transfected with the gRNA1, gRNA2 or scramble along with the donor cassette using TurboFectin (Origene, cat# TF81001). Forty-eight hours post-transfection, cells were split 1:10. Cells were split 1:10 a total of seven times. Transfected cells were selected for using puromycin at 8 ug/mL. Monoclonal colonies were established using cloning discs soaked in trypsin. Viable colonies were grown to confluence and tested for heterozygous or homozygous knockout of BMAL1 using PCR and western blot analysis.

PCR primers used to confirm genomic integration were as follows for the left integration junction: TACTAATGTAGCCCAGGATGGT (Sense) and TAGGTGCCGAAGTGGTAGAA (AntiSense); right integration junction: AATGGAAGGATTGGAGCTACG (Sense) and CTCAATGATCTGGGATGACTTACA (AntiSense); and donor cassette CAGATGCCGGTGAAGAAAGA (Sense) and GGAATGAGCTGGCCCTTAAT (AntiSense). PCR amplified DNA was Sanger sequenced at Purdue University’s Genomics Core.

Preliminary growth curve analysis studies indicated multiple monoclonal lines created using scramble sequences exhibited off target effects on cell growth. Blasting the scramble sequence against the mouse genome found multiple growth regulatory genes near similar sequences. Alterations in coding sequence, etc., due to integration of scramble could have affected these genes. Thus, a negative control was abandoned for all subsequent studies.

### Fluorescence activated cell sorting (FACS) analysis of percent of proliferating cells

For FACS analysis of percent proliferating cells across eight-days of culture, cells were plated at a density of 100,000 cells/ml, and duplicate samples of each line were harvested on days 2, 4 and 8. Cells were counted and approximately 1.5 x10^6^ cells were pelleted by centrifugation. Cell pellets of were resuspended in 100 µl of PBS, fixed with drop-wise addition of 280 µl of ice-cold 90% ethanol and stored at −20° C until day of analysis. On day of analysis, fixed cells were pelleted by centrifugation and resuspended in 1ml PBS-0.5% BSA, pelleted again, resuspended in PBS-0.5% BSA with 100 U/ml RNAse (R6513 Sigma-Aldrich), and incubated for 15 min at 37°C. Five µl of propidium iodide (PI; 500 µg/ml stock solution) was added and cells were incubated 15 minutes at RT. PI labelled cells were analysed by fluorescently activated cell sorting (FACS) using a Beckman Coulter FC500 instrument in the Purdue University’s Flow Cytometry and Cell Separation Facility. FACS analysis was repeated 3-times.

### MTT cell proliferation and reactive oxygen species (ROS) assays

HC11 and BMAL1-KO cells were plated in 96-well plates at 3,000 cells/well in RPMI 1640-phenol free growth media. For MTT (3-(4,5-dimethylthiazol-2-yl)-2,5-diphenyltetrazolium bromide) assay, on day 2, 4, 6, and 8 after plating (three wells per line per day), media was removed from wells and cells were incubated with MTT from the Cell Proliferation Kit (Sigma Aldrich, catalogue no. 11465007001), and the colorimetric assay was performed following manufacturer’s directions. After completion of the assay, images of cells were captured under bright field at 10 X magnification with a Nikon camera attached to a Zeiss inverted microscope, and absorbance was read at 540 nm on the Spark multimode microplate reader (Tecan Trading AG, Switzerland). To assess relative amount of activity per cell on days 2 and 6 of culture, MTT staining per unit area of cells was measured using the ColonyArea Plugin [61] to capture and calculate colony intensity percent. Colony intensity percent is the ratio of the sum of pixel intensities in a region to the sum of all the pixels within the same region of interest multiplied by 255, i.e. assuming highest intensity with full saturation of these pixels. For analysis 100 individual cells were selected across three images and percent intensity of staining was measured. The Reactive Oxygen Species Assay Kit (Abcam catalogue No. ab113851) was used to measure the level of ROS in cell lines on days 3 and 4 (three wells per line per day) after plating cells, with fluorescence immediately measured at Ex/Em 485/535 nm on the Spark multimode microplate reader. Both assays were performed three times.

### Protein Isolation, Enzyme-linked Immunosorbent Assay (ELISA), Western Blot and Immunoprecipitation (IP) Analysis

Protein lysates were isolated from cultures by pouring off media, washing twice with chilled PBS, and harvesting using a scraper and 3 ml of cold PBS. Cells were pelleted by centrifugation, and cell pellets were lysed for 30 min on ice with 600 µl of Cell Extraction Buffer (Invitrogen, supplemented with 1mM of PMSF and 50 µl/ml of Protease Inhibitor Cocktail, Sigma Aldrich), with vortexing at 10 min intervals. Protein lysates were transferred to microcentrifuge tubes and centrifuged at 13,000 rpm for 10 min at 4 ° C. Protein concentration was measured using a nanodrop (ThermoFischer) and the Coomassie Plus Protein Assay kit (Pierce Coomassie Plus (Bradford) Protein Assay; Thermo Fisher Scientific) following the manufacturer’s protocol. Samples were stored at −80°C until further analysis.

BMAL1 (cat. no. LS-F39618) and beta-casein (CSN2, cat. no. LS-F13103) protein levels in cell lysates were measured with ELISA kits from LSBio following manufacturer’s directions. Since number of cells and protein concentration changed with state of differentiation of cells, data were expressed as micrograms of CSN2 or BMAL1 protein per mg of protein.

For western blot analysis, 100 µg of protein were loaded per lane and electrophoresed on a 10% TGX precast SDS PAGE gel from Bio-Rad. Protein was transferred onto a nitrocellulose membrane, and membranes were blocked using either 5% BSA (Bovine Serum Albumin), and probed for BMAL1, CSN2 and beta-actin proteins using anti-BMAL1 (ab3350, dilution 1:1,000), anti-CSN2 (LS-C373659, Life Spam, dilution 1:200) and Anti-beta-actin (AbCam ab8227, dilution 1:10,000) antibodies, respectively. Blots were washed and then incubated with secondary antibody (ab97051; 1:5000). Membranes were washed and incubated with the detection reagent Clarity Western ECL Substrate (Bio-Rad). Blots were imaged using the ChemiDoc MP system (BioRad). ImageJ was used for densitometric analysis. Density of test protein (BMAL1 and CSN2) were divided by density of beta-actin (BA) band. To enable statistical analysis across multiple gels run to measure CSN2 protein levels, ratio was normalized to relative expression of HC11 UNDIFF.

For IP analysis of specificity of ChIP grade antibody BMAL1 (Abcam; ab3350), rabbit polyclonal antibody to BMAL1 (Abcam; ab3350) was added to 200 µg protein in 200 µl cold PBST (PBS pH 7.4 with 0.02% Tween-20), and rotated overnight at 4° C. The antibody protein mixture was added to Dynabeads protein A for immunoprecipitation (ThermoFischer) and pipetted gently to resuspend. Samples were rotated for 2 hr at 4° C. Tubes were placed on magnetic rack to pellet beads, and washed three times with 200 µl cold PBS. Then 15 µl PBS and 15 µl 2X Laemmli sample buffer with 2-mercaptoethanol added per manufacturer’s instructions, and gently pipetted to resuspend. Samples were heat at 100° C for 5 min and placed on magnetic rack to pellet beads. Supernatant was removed and separated using SDS-Page gel for analysis by western blot. Mouse monoclonal BMAL1 antibody (Santa Cruz Biotechnology, Inc., Dallas, TX US; cat. no. sc-365645) was used for western blot analysis.

### RNA Isolation and Real-Time Quantitative PCR Analysis (RT-qPCR)

RNA was collected from cells and isolated using Qiagen’s RNeasy kit with an on-column DNase treatment, and quantity was measured using Nanodrop. RIN scores of total RNA following nanochip analysis on an Agilent 2100 Bioanalyzer were found to vary from 8.0-9.0 across all samples. Promega’s GoScript Reverse Transcriptase kit was used to reverse transcribe 500 ng of total RNA into cDNA following manufacturer’s protocol. Gene expression was analysed with TaqMan Assays on Demand using 2X TaqMan Gene Expression Master Mix (Life Technologies) using a 1:10 dilution of the cDNA product. The CFX Connect Real-Time PCR Detection System (BioRad) was used to run RTq-PCR, which was initiated with 2 min incubation at 95°C and then 40 cycles of 95°C for 15 sec and 60°C for 1min, and standby was set at 4°C. Multiple genes (beta-actin, beta microglobulin and 18S) were screened as reference genes (housekeeping gene) for calculating relative expression using the 2^-ΔΔ CT^ method [62]; 18S was chosen as the reference gene based on its levels staying steady across time and genotype. Mouse-specific Assay on Demand TaqMan assays (Life Technologies) used were *Per2* (Mm00478099_m1), *Sod3* (Mm01213380_s1), *Csn2* (Mm04207885_m1), *Tph1* (Mm01202614_m1), *Slc6a4* (SERT, Mm00439391_m1), *Ccnd1* (Mm00432359_m1)*, Ppara* (Mm00440939_m1), *Prlr* (Mm04336676_m1), and *18S*. To calculate relative expression, mean ΔCT of wild-type HC11 UNDIFF cultures was used as normalizer for relative expression. For temporal analysis of *Per2* expression across 48 hr sampling mean ΔCT of HC11 cultures across all time was used as the normalizer for relative expression. Data are presented as mean fold-change ± standard deviation of relative expression levels, or mean of log base two-fold change ± standard deviation for *Csn2* expression.

### ChIP assay and q-PCR verification of known targets

Use of two independent biological replicates for ChIP-seq analysis to identify transcriptional targets is consistent with ENCODE (Encyclopaedia of DNA Elements) guidelines[63]. For ChIP-seq studies, we performed four independent ChIP experiments, two with undifferentiated cultures and two for differentiated cultures, to enable analysis of the effect of differentiation on BMAL1 targets. Briefly, undifferentiated and differentiated HC11 cultures in 100 mm plates were treated with 1% formaldehyde to cross-link proteins to chromatin by adding 625 µl fresh 16% formaldehyde directly to the dishes containing cells and 10 mL media, and incubated at RT on a shaking platform for 7 min. To quench the formaldehyde, 500 µl of 2.5M glycine was added, and incubated at RT on the shaking platform for 5 min. Cells were washed twice with ice cold PBS, and removed from culture dish using a cell scraper. Cells were pelleted by centrifugation and stored at the −80 °C. Samples were collected for ChIP assays in four replicate experiments (sample IDs UNDIFF1, UNDIFF2, DIFF1, DIFF2).

Nuclei were collected by resuspending each fixed-cell pellet in 10 mL of Rinse 1 (50mM Hepes pH 8.0, 140mM NaCl, 1mM EDTA, 10% glycerol, .5% NP40, .25% Triton x100), and incubating 10 min on ice. Samples were pelleted by centrifugation 1,200 X g at 4 °C for 5 min, and then resuspend in 10 mL CiA NP-Rinse 2 (10mM Tris pH 8.0, 1mM EDTA, 0.5mM EGTA, 200mM NaCl), and then centrifuged at 1,200 X g and 4 °C for 5 min. Pellets were washed with 5 ml of CiA Covaris Shearing Buffer (0.1% SDS, 1mM EDTA pH 8.0, 10mM Tris HC1 pH 8.0) twice with centrifugation at 1,200 X g and 4 °C for 3 min after each wash. For sonication, pellets from approximately 3 million cells each were resuspended in 130 µl of CiA Covaris Shearing Buffer with 10% protease inhibitor cocktail. Pellets were sonicated in a Covaris E210 instrument (Covaris, Inc., Woburn, MA) using the following parameters: Duty Factor 5%, Peak power 105W, Cycles per burst 200, 10 min. Following sonication sheared chromatin lysate was transferred to a new centrifuged at 20,000 X g and 4 °C for 15 min. Supernatant (sheared chromatin stock) was divided into three aliquots for input DNA, mock-immunoprecipitation (IP) and IP and stored at −80 °C (−20C), after removal of an aliquot for analysis of product size in a 2% agarose gel and on the Agilent Bioanalyzer Nanochip.

For immunoprecipitation step, cross-linked chromatin using anti-BMAL1 antibody and mock-IP aliquots were thawed on ice. Samples were precleared by adding 50 µL resuspended A beads (DynaBeads Protein A, Invitrogen cat. # 10001D), and incubating at 4° C for 1.5 hr with rotation. Tubes were placed on a magnetic rack and liquid was transferred to a new microfuge tube. One-quarter of the volume of 5X IP Buffer (250mM Hepes/KOH pH 7.5, 1.5M NaCl, 5mM EDTA, 5% TritonX 100, 0.5% DOC [4-chloro-2,5-dimethoxy-amphetamine], 0.5% SDS) was added, and 10 µL of BMAL1 antibody (Abcam ab 3350) was added to IP tube, and nothing was added to mock-IP tube. Samples were rotated overnight at 4° C, and then 50 µL of resuspended A beads were added to each tube, and rotated for 3 hr at 4° C. Magnetic rack was used to remove A beads. Supernatant was collected transferred to a new tube, and washed with 1mL IP Buffer twice with rotation at RT for 3 min, and collection on magnetic rack following wash. Supernatants were then washed with 1 mL of DOC buffer (10 mM Tris, pH8.0; 0.25 M l (iCl; 0.5% NP40, 0.5% DOC, 1 mM EDTA), followed by 1 mL of Tris EDTA (TE) buffer pH 8. Supernatant was removed and 150 µL elution buffer was added and samples were rotated at RT for 20 min. Supernatant was transferred to a new tube and store at −80 °C until DNA isolation.

Chromatin was isolated from supernatant by adding 3 volumes of TE with 1% SDS and RNAse A (Sigma R6513), and vortexing to mix. Samples were incubated at 37° C for 30 min, and proteinase K was added, vortexed again to mix, and cross-linking was reversed by incubating at 55° C for ∼2.5 hr, and then overnight at 65° C in PCR tubes in the thermocycler. DNA was extracted the next day by adding 230 µL TE and 330 µL phenol/chloroform, vortexing, followed by centrifugation for 1 min. Aqueous layer was transferred into a new microfuge tube and 10 µL of commercial glycogen (ThermoScientific R0551) was added with 30 µL 3M NaCl. One volume of 100% ethanol was added and samples vortexed to mix. Samples were incubated for two days at −20 ° C, and then centrifuged 1 hr at 4 °C to pellet DNA. Pellet was washed twice by removing the supernatant and adding 500 µl 70% EtOH. Pellet was resuspended in 25 µL TE and concentration was measured with Qubit fluorometer (Thermo Fisher Scientific) and a Bioanalyzer 2100 (Agilent), and samples stored at −80° C until sequencing.

To determine specificity BMAL1 antibody, qPCR analysis of ChIP product, input and mock-IP samples were performed using primers designed to target the promoter region of *Per1*, a transcriptional target of BMAL1: CLOCK, and the exon region of a mouse sperm gene *Magea1_2*, which was not expected to be a target of BMAL1 binding. The promoter region of *Per1* containing the proximal E-box enhancer was amplified with the following primer set: forward, 5-CCTCCCTGAAAAGGGGTA-3; reverse, 5-GGATCTCTTCCTGGCATCTG-3. Primers used to amplify MAGEA1_2, with forward: 5-GCCTCTGAGTGCTTGAAGAT-3, and reverse: 5-CAGGGCAGTGACAAGGATATAG-3. Primers were designed to promoter regions of SLC6A4 (aka SERT) to encompass an E-box most proximal (beginning at −42 nucleotides) to the transition start site forward: 5-CTCCAGCTGCGGTAGCAGA-3 and reverse: 5-ATTTGTACTTGCGGCCC-3, and more distal (beginning at −1282 nucleotides) Forward: 5-GGAGTTACAGGCACGGAAG-3 and reverse 5-GCCTGGCCATTCCATGA-3. Triplicate samples were measured using SYBR green real-time PCR master mix (ThermoFischer Scientific). The analysis was conducted using the CFX Connect Realtime PCR Detection System (Biorad) with the following conditions for reactions were 1 cycle at 50°C for 2 min; 1 cycle at 95°C for 2 min, 55 cycles at 95°C for 15 sec, gradient 58-60°C for 30 sec and 72 °C for 1min; and 1 cycle at 95°C for 10 sec, melt curve 65°C – 95°C, increment 0.5°C for 5 sec. PCR reaction efficiencies were 98% at annealing temperature 58°C for *Per1* and *Sert*, and at 60°C the efficiency for *Magea1* was 96%.

To calculate relative amount of input target brought down by ChIP with BMAL1 antibody the following equation was use: adjusted input sample-C_T_ of ChIP sample (https://www.thermofisher.com/us/en/home/life-science/epigenetics-noncoding-rna-research/chromatin-remodeling/chromatin-immunoprecipitation-chip/chip-analysis.html). A positive ChIP was defined as at least 2-fold greater than mock-IP sample [64].

### Sequencing of DNA, peak identification, and functional annotation analysis

Input and ChIP sample pairs were sequenced together by paired ends reads on the Illumina Novaseq 6000 (San Diego, CA) following the manufacturer’s protocols. Prior to sequencing libraries were prepared using NEXTFLEX Rapid DNA-Seq Library Prep Kit for Illumina Platforms (PerkinElmer, Waltham, MA) according to manufacturer’s protocol. The number of reads for three of the input ChIP sample pairs (both UNDIFF samples and one DIFF sample) averaged approximately 100 million reads. The remaining DIFF input-ChIP pair had over 1 billion reads. To account for discrepancy in depth of sequencing, the reads were divided into five groups, and one group of input-ChIP sample pair was used for analysis. Sequence quality was assessed using FastQC (v 0.11.7) for all samples and quality trimming was done using Fastx toolkit to remove bases with Phred33 score of less than 30. The resulting reads of length at least 50 bases were retained. The quality trimmed reads were mapped against the reference genome (Mus_musculus.GRCm38.chr.fa) using bowtie2 (Version 2.2.9)[65]. ChIP-seq data were submitted to Gene Expression Omnibus (GEO accession number GSE154937).

Peaks were called from mapped files (bam files) using ‘callpeaks’ script of HOMER (Version 4.10)[66] to detect transcription factor associated peaks using default settings except for the following parameters: fold enrichment over input tag count (F) was set at 2, fold enrichment limit of expected unique tag positions (C) was set at 2 and fold enrichment over local tag count (L) was set at 2 and FDR was set at <0.01. Quality control analyses were run from files generated in read tag directories to include tag count distribution, autocorrelation analysis, genomic nucleotide frequency relative to read positions, and fragment GC % distribution, met criteria for further analysis. Picard Mark Duplicates was performed on each sample (Supplemental Table S18). Common nearest gene associated with peaks between UNDIFF1 and UNDIFF2, DIFF1 and DIFF2 were generated using Venny 2.1[67].

Overlap of BMAL1 targets in HC11 cells with genes that exhibit circadian rhythms of gene expression in virgin mouse mammary glands[13] and in human breasts during lactation[14] was investigated by obtaining supplemental data from these manuscripts. Similarly, overlap of HC11 BMAL1 targets with BMAL1 transcriptional targets identified in mouse liver was also investigated by obtaining supplemental data this manuscript[19]. For comparative analysis, gene symbols were converted to mouse ENSEMBL IDs using the gene conversion tool available in DAVID, and then common genes across data sets were identified using Venny 2.1[67].

Functional annotation analysis was performed using Database for Annotation, Visualization, and Integrated Discovery-DAVID Bioinformatic Resources 6.8[68, 69], and Ingenuity Pathway Analysis (IPA; QIAGEN Inc., https://www.qiagenbioinformatics.com/products/ingenuitypathway-analysis). To aid in visualization of IPA generated figures, genes were assigned a value of 5 and were indicated as red in figure of Regulator Networks. The network with highest consistency score was used, with the higher scores reflecting paths between target gene and output function consistent with the predicted state of the regulator—here all indicated as upregulated--based on the literature. IPA predicted upstream regulators were removed from the generated network, as BMAL1 was the upstream regulator (Supplemental Figure S2). The My Pathway tools in IPA was used for visualization of a subset of BMAL1 target genes. GeneCards [70] was used to query information on function of genes of interest.

### Statistical analysis

Statistical analysis was performed using SPSS software (IBM SPSS, v.26). General linear model was used to analyze whether cell line (HC11 or BMAL1-KO) or state of differentiation (DIFF or UNDIFF), or day (growth curve, MTT assay, ROS assay, and FACS analysis) significantly impacted variables (relative expression, protein abundance, doubling time, and cell density). A Tukey’s post-hoc test was used for pairwise comparisons. Significance was considered at P ≤ 0.05. Cosine fit analysis of 24 hr rhythms of gene expression was performed with the cosinor package in R (RStudio 1.1.453, Boston, MA). Mesor, amplitude, acrophase R^2^, and *p*-value were outputs of the package algorithm.

## Acknowledgements

The authors would like to thank Paul Parker for analysis of ChIP samples, and Shaojun Xie for reading and editing manuscript.

## Authors’ contributions

TC: conceived ideas, designed studies, analyzed data, wrote manuscript; AT: conceived ideas, designed and conducted studies, analyzed data, edited and approved manuscript, SC: designed and conducted studies, analyzed data, edited and approved manuscript; KH: designed and conducted studies, analyzed data, edited and approved manuscript; JC: designed and conducted studies, analyzed data, edited and approved manuscript; KB designed and conducted studies, analyzed data, edited and approved manuscript; CA: designed and conducted studies, analyzed data, edited and approved manuscript, KT: conducted studies, analyzed data, edited and approved manuscript; AS: conceived ideas, designed studies, edited and approved manuscript; SM: conceived ideas, designed studies, edited and approved manuscript; PSM: designed and conducted studies, analyzed data, edited and approved manuscript; JT: designed and conducted studies, analyzed data, edited and approved manuscript, KP: conceived ideas, designed studies, edited and approved manuscript. All authors approved the manuscript.

## Data availability statement

ChIP-seq were made publicly available through Gene Expression Omnibus (GEO) and can be found using the following accession number GSE154937. All other data will be made available upon request to corresponding author.

---

**Supplemental Figure S1.** (a) Western blot analysis of immunoprecipitation (IP) of BMAL1 protein in HC11 cell protein lysate using rabbit polyclonal antibody to BMAL1 (ab3350, ChIP grade, 2 µg per IP). Lane 1 = Precision Plus Protein Standard Ladder; Lane 2= 100 µg, Lane 3= 150 µg, Lane 4 = 200 µg, Lane 5= 200 µg of lysate precleared and incubated with beads but no Ab (-), Lane 6= an aliquot of supernatant pulled off of lane 5 sample before wash steps that precede elution. western blot was performed using the mouse monoclonal antibody to BMAL 1 (sc-373955 @ 1:750 primary antibody concentration) for visualization. (b) Electropherogram analysis of input DNA used for ChIP-seq shows ideal size for next generation sequencing (seq). (c) Evaluation of antibody specificity indicated no difference between mock-ChIP and BMAL1-ChIP samples in the cycle threshold values following q-PCR analysis of an exon region of the *Magea1_2* sperm specific gene, which is not a BMAL1:CLOCK target. Whereas a 9-fold difference in enrichment was found between q-PCR analysis of BMAL1-ChIP and mock-ChIP for the *Per1* promoter region versus the exon region of *Magea1_2.* To calculate relative amount of input target brought down by ChIP with BMAL1 antibody the following equation was use: adjusted input sample-C_T_ of ChIP sample. A positive ChIP was defined as at least 2-fold greater than mock-IP sample.

**Supplemental Figure S2.** Ingenuity pathway analysis generated network with greatest consistency score reflecting relationships between BMAL1 targets and predicted downstream effects. Gene names and symbols are defined in Supplemental Table S13. The predicted upstream regulators were removed, as analysis assumes this is BMAL1 as the transcriptional regulator of genes in red and combined the downstream effects of inhibiting organismal death, and stimulating efflux of lipids, cellular protrusions, branching of cells, development of sensory organ, sprouting and organ size.

**Supplemental Figure S3.** (a) PCR Analysis of monoclonal donor cassette integration. The first 10 lanes contain DNA from monoclonal colonies post clonal selection. The first character of each name refers to the gRNA used (i.e. 1A is HC11 that has undergone integration using gRNA 1). NTC denotes a no-template negative control. These results indicate proper integration of the donor cassette into the target site of cell colonies 1B, 1C, 2A, 2B, 2C, and 2D.

**Supplemental Figure S4.** Figures represent reads/samples in wig format where Input reads are subtracted from IP reads. Reads per sample of peaks closest to: *Ppara* zoomed out (a) and in (b); *Prlr* (c) and (d); *Fasn* (e); *Slc6a4* (f); *Per1* (g); and *Csn2* (h) transcriptional start sites.

**Supplemental Figure S5.** Quantitative-PCR (q-PCR) analysis of *Fasn* of HC11 (black) and BMAL1-KO (gray) cultures in replicate experiments of undifferentiated (UNDIFF) and lactogen differentiated (DIFF) cultures. Values are mean technical replicates within experiment, normalized to express fold-change relative to mean of HC11 ± standard deviation using delta-delta cycle threshold method. Temporal analysis of *Fasn* expression in WT HC11 (solid line) and BMAL1-KO (dashed line) cultures. For this experiment cells were grown to confluence in growth media. Media was changed to lactogen media for 2 hr to synchronize clocks. At completion of 2 hr lactogen treatment (time 0 hr), cells were rinsed with PBS and cultured in growth media for remainder of the experiment. Cells were collected for isolation of total RNA every 4 hr over a 48 hr period beginning at 0 hr. *Fasn* was measured with q-PCR, and levels were expressed relative to mean levels across all time points of HC11 culture. Cosinor analysis found mesor (0.01 and 0.97), amplitude (0.21 and 0.12), acrophase (−9.14 and −4.44), R^2^ (0.26 and 0.19) and *p*-value (0.22 and 0.33) of fit to a 24 hr rhythm were calculated, respectively, for HC11 and BMAL1-KO lines.

